# Evolutionary rewiring of the DNM2 proline-rich domain drives lineage-specific constraints in erythropoiesis

**DOI:** 10.64898/2026.03.20.713161

**Authors:** Thomas Arbogast, Peggy Kirstetter, Christine Kretz, Claire Masson, Susan Chan, Philippe Kastner, Anita Eckly, Jocelyn Laporte

## Abstract

Dynamin-2 (DNM2) is a ubiquitously expressed GTPase essential for membrane trafficking, yet how its evolution shapes tissue-specific functions remains unclear. Here, we generated a humanized mouse model in which murine *Dnm2* is fully replaced by human *DNM2*. Human DNM2 rescues the embryonic lethality of *Dnm2* deletion but cannot sustain postnatal hematopoiesis, leading to fatal hemolytic anemia, erythroblast maturation arrest, reticulocytosis, and macrothrombocytosis. The defect emerges selectively during postnatal bone marrow erythropoiesis and is associated with impaired transferrin receptor turnover, despite normal fetal liver hematopoiesis. Comparative proteomics reveals extensive species-specific rewiring of the DNM2 proline-rich domain (PRD) interactome, with GRB2 as the sole conserved partner. *Dnm2* haploinsufficiency phenocopies the erythroid defect with reduced severity, indicating a dosage-sensitive loss-of-function mechanism. These findings identify PRDs as rapidly evolving context-dependent adaptors that tune ubiquitous proteins to tissue-specific demands, and highlight regulatory domains compatibility as a key consideration for humanized models and gene-replacement strategies.

## INTRODUCTION

The dynamin superfamily consists of large GTPases which play a crucial role in membrane tubulation, scission and fusion by oligomerizing at the neck of membrane invaginations along membrane structures^1–3^. In addition to their role in membrane remodeling and trafficking, dynamins are involved in cytoskeleton rearrangement^4^ which influences both cell migration^5^, and cell division^6^. Structurally, classical dynamins are composed of several functional domains including a N-terminal GTPase domain, a middle domain or stalk domain involved in protein oligomerization, a GTPase effector domain (GED), a pleckstrin homology (PH) domain that interacts with phosphoinositides. Additionally, they include a C-terminal proline-rich domain (PRD) that binds SH3 domain-containing proteins including amphiphysins, endophilins and syndapins^7^. Dynamin forms tetramers in solution and the auto-inhibition driven by the interaction between the stalk and the PH domains is released upon lipid binding, allowing the protein to adopt an open configuration^8,9^. This conformational change enables dynamins to assemble into a helical structure, which stimulates GTP hydrolysis concomitant to membrane constriction and fission. In mammals, three classical dynamins are encoded by *DNM1*, *DNM2*, and *DNM3* and exhibit approximately 80% amino acid sequence identity, with the PRD domain showing the most divergence with 60-70% identity. While *DNM1* and *DNM3* are predominantly expressed in neurons, *DNM2* is ubiquitously expressed across tissues, with its highest expression reported by the GTEx database observed in whole blood in Human.

In contrast to this wide expression pattern, disease-causing dominant mutations in *DNM2* have been associated to three autosomal dominant neuromuscular diseases: centronuclear myopathy^10^ (CNM), Charcot-Marie-Tooth disease^11,12^ (CMT), and hereditary spastic paraplegia^13^ (HSP). Mutations linked to CNM and CMT cluster in the middle and PH domains^14^ while the mutation linked to HSP is localized in the GED domain^13^. Although the precise pathogenic mechanisms underlying these conditions are unclear, an imbalance of DNM2 activity appears to play a critical role. Specifically, gain-of-function mutations have been associated with CNM, whereas loss-of-function mutations have been associated with CMT respectively^15–18^. Beyond neuromuscular disorders, mutations in *DNM2* have been found in cancers^19^ including T-cell acute lymphoblastic leukemia^20^, and linked to neutropenia^21–23^. These findings suggest the importance of DNM2 in the development and maintenance of hematopoietic cells.

To study the tissue-specific associated diseases, several mouse models carrying *Dnm2* mutations associated with CNM and CMT have been generated and characterized. Notably, while mouse models carrying CNM mutations successfully replicate the phenotypes observed in patients^24–26^, a mouse model carrying the K562E mutation that causes CMT in humans exhibit characteristics of a primary myopathy^27^. These findings suggest potential differences in tissue-specific *DNM2* expression, interactome composition, and/or function between humans and mice.

Because the physiological role of human DNM2 and its tissue-specific functions remain insufficiently defined, thereby limiting the translational relevance of murine models for human disease, we generated a humanized (Hum) mouse model lacking endogenous murine *Dnm2* and expressing human *DNM2*. The present study integrated multiscale phenotyping across tissue, cellular, ultrastructural, proteomic, and comparative evolution levels. While constitutive knock-out of *Dnm2* is embryonic lethal^28,29^, Hum mice are born but rapidly develop hemolytic anemia, poikilocytosis, and macrothrombocytosis, in the absence of detectable neuromuscular abnormalities. Comparative analysis of the human and mouse PRD interactomes revealed marked species-specific differences, uncovering a previously unrecognized species-specific adaptation of dynamin 2, particularly within hematopoietic lineages.

## RESULTS

### Generation and validation of a DNM2-humanized mouse model

To tackle the physiological role of human DNM2 and its tissue-specific functions, we generated *DNM2*-humanized (Hum) mice in which endogenous murine *Dnm2* genes are ablated and replaced by two copies of the human DNM2 gene from a human bacterial artificial chromosome (BAC) transgene (Tg; Supplementary Fig. 1). Crosses between *Dnm2^+/-^*and Tg(*DNM2*)_*Dnm2^+/-^* mice produced offspring with expected genotypic *Dnm2* ratios (25.5% +/+, 49.5% +/- and 25% -/- over 204 males), suggesting no prenatal lethality in Hum and Tg mice (Supplementary Fig. 2A). The *DNM2* transgenes being located on the Y chromosome, only males were used for this study. Wild-type (WT) animals were generated separately by mating *Dnm2^+/-^* mice together. We validated the absence of endogenous *Dnm2* expression in Hum mice and the absence of *DNM2* transgene expression in WT mice at both RNA and protein levels (Supplementary Fig. 2B, C, F, G, 10A-F & Table 1). We further verified that the human transgene was uniformly expressed across organs of Hum mice (Supplementary Fig. 2H & 10G-I). Expression of other dynamin paralogs revealed partial genetic compensation, with downregulation of *Dnm1* and a five-fold upregulation of *Dnm3* relative to WT littermates (Supplementary Fig. 2D, E). Overall, Hum mice establish a robust platform to interrogate species-specific DNM2 function *in vivo* (Fig. 1A).

**Figure 1.**
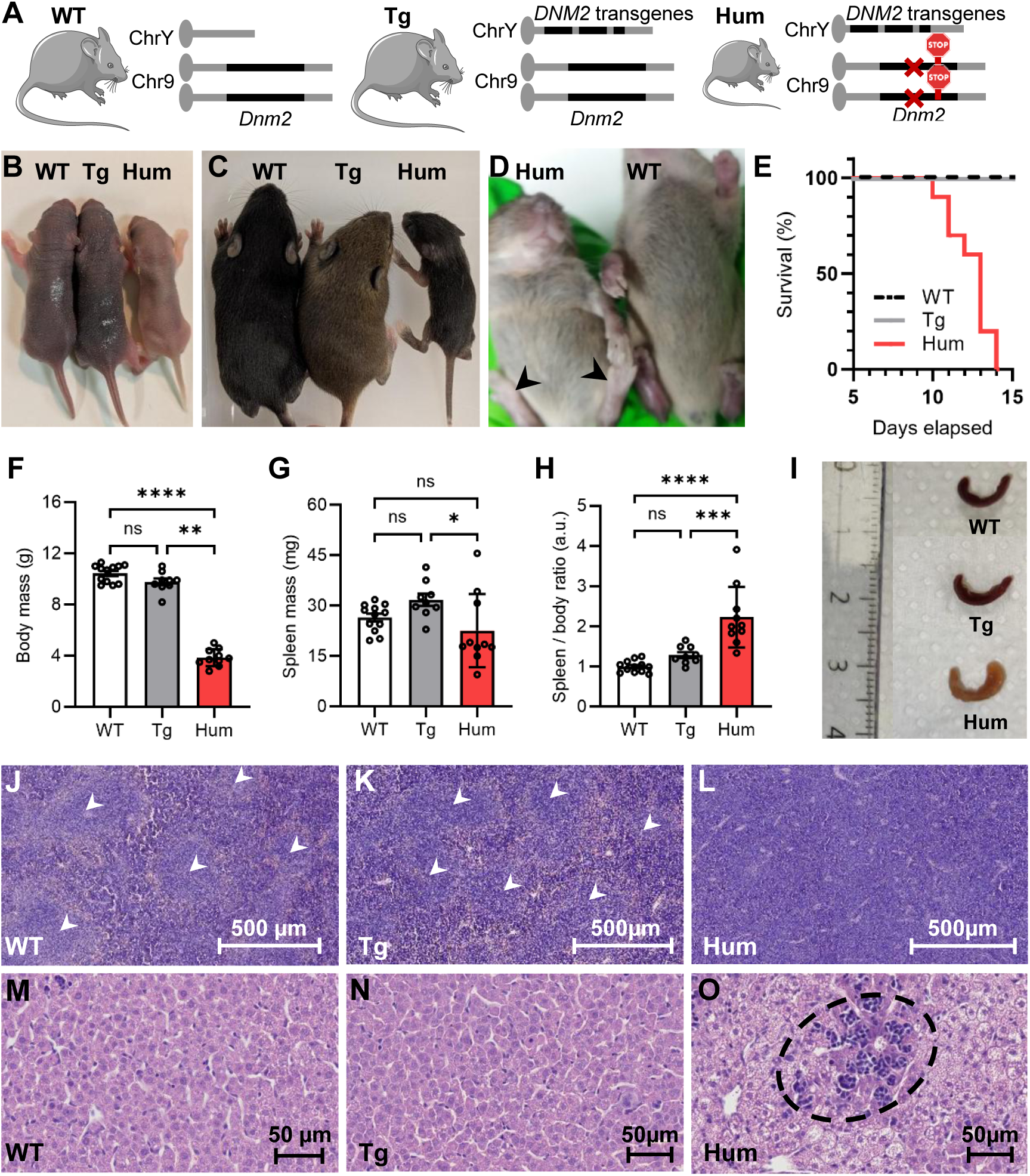
Humanized mice have limited viability, show splenomegaly and extramedullary hematopoiesis. (**A**) Genetic definition of the transgenic (Tg) and humanized (Hum) mouse models used in this study. **(B, C**) Dorsal view of WT, Tg, and Hum littermates at P3 and P12. (**D**) Ventral view of WT and Hum littermates at P12. Black arrowheads indicate pale-color hindlimb extremities of Hum mice. (**E**) Viability of animals. **(F)** Body mass at P12 (g). (**G**) Total spleen mass at P12 (mg). (**H**) Spleen/body ratio (arbitrary unit). (**I**) Lateral view of whole spleens. (**J**-**O**) Histological characterization. (**J**-**L**) Longitudinal spleen sections at low magnification. Alternation of white pulp (white arrowheads) and red pulp were observed in WT and Tg sections whereas only red pulp was observed in Hum sections. (**M**-**O**) Longitudinal liver sections at high magnification. The dashed circle indicates extramedullary hematopoiesis. Steatosis was observed homogenously in Hum liver sections. Data are mean ± SD, ns: not significant, \**P* < 0.05, \*\**P* < 0.01, \*\*\**P* < 0.001, \*\*\*\**P* < 0.0001. (**G**, **H**) One-way ANOVA + Tukey’s post-hoc; (**F**) Kruskal-Wallis + Dunn’s post-hoc.

### DNM2-humanized mice exhibit postnatal growth failure and early lethality

While expression of human DNM2 rescued the embryonic lethality of *Dnm2^-/-^* mice, Hum animals displayed growth delays shortly after birth, and were consistently smaller and lighter than the other genotypes at P12 (WT, 10.4g; Tg, 9.8g; Hum 3.9g; Fig. 1B, C, F; Supplementary Table 2). Additionally, Hum mice exhibited impaired mobility upon stimulation and pale paws (Fig. 1D). All Hum animals died between P11 and P14, whereas Tg and WT animals remained healthy (Fig. 1E). These findings indicate that human DNM2 cannot fully support essential postnatal physiological functions normally mediated by murine Dnm2.

### Lethality of humanized mice is associated with splenomegaly and extramedullary hematopoiesis

To decipher the species-specific role of DNM2 in physiology, Hum animals were euthanized at P12 and the underlying cause of their mortality was investigated. The autopsy of Hum mice revealed markedly pale-colored internal organs and peripheral blood compared to WT and Tg littermates, indicating anemia (Supplementary Fig. 3). Consequently, we first evaluated the spleen of animals which was pale and hypertrophied in Hum mice (Fig. 1G-I). Unlike humans, the rodent spleen is predominantly composed of white pulp which is implicated in adaptative immunity. Histological analysis revealed that while WT and Tg spleens displayed normal alternation of white and red pulps, Hum spleens showed only red pulp (Fig. 1J-L) which serves as the primary site for the degradation of senescent erythrocytes^30,31^ and, in rodents, functions as the major zone of extramedullary hematopoiesis^32,33^. This anomaly, coupled with splenomegaly, suggests significant disruptions in red blood cell (RBC) production and/or degradation in Hum animals. While WT and Tg animals had normal liver histology and hepatocytes, livers of Hum mice exhibited centrilobular microvesicular steatosis, a histological marker of anemia-induced hepatic stress (Fig. 1M-O). Extended extramedullary hematopoiesis was also evident in the liver of Hum mice, likely secondary to splenic hematopoiesis. These changes are consistent with severe hematological stress and compensatory erythropoiesis, suggesting that failure of the hematopoietic system is a primary driver of postnatal lethality.

Given that *DNM2* mutations are linked to centronuclear myopathy (CNM) and dominant intermediate Charcot-Marie-Tooth (DI-CMT) diseases, we also assessed sciatic nerve and gastrocnemius muscle structures (Supplementary Fig. 4). Coronal sections of the sciatic nerve showed no myelination defects nor fiber degeneration in both Hum and Tg mice. Gastrocnemius muscle coronal sections revealed no central nuclei or fiber size abnormalities in Hum and Tg littermates. The ultrastructure analysis by electronic microscopy confirmed normal Z-line, triads, and mitochondria organization. In skeletal muscle, one major isoform incorporates the alternative exon 12b–referred to as *M-DNM2* (muscle-specific)–while the isoform lacking this exon is known as *Ub-DNM2* (ubiquitous)^34^. RT-qPCR analysis of liver and quadriceps samples showed that the human transgene expresses both *Ub-DNM2* and *M-DNM2* in Hum quadriceps, whereas only the *Ub-DNM2* isoform is detected in Hum livers (Supplementary Fig. 4G, H & Table 1). These findings suggest functional equivalence of mouse and human DNM2 in muscle and nerve tissues, although the progressive nature of CNM and CMT may postpone the appearance of muscle and nerve abnormalities beyond the timeframe of this study.

### Humanized mice develop severe hemolytic anemia with macrothrombocytosis

Histological evidence strongly suggests that Hum mice experienced anemia, which prompted compensatory erythropoiesis in both the spleen and liver. To confirm this hypothesis, we conducted hematological analyses (Fig. 2, Supplementary Tables 3 & 4). In comparison to WT and Tg, Hum mice exhibited a five-fold decrease in RBC count and hemoglobin (WT, 11.2g/dL; Tg, 12.3g/dL; Hum 2.4g/dL), a decrease in mean corpuscular volume, and a six-fold decrease of hematocrit (Fig. 2A-D), all of which are indicative of severe anemia. Elevated red blood cell distribution width (RDW) showed anisocytosis, and peripheral blood (PB) smears revealed poikilocytosis, hemolysis, and hypochromia (Fig. 2E). Hum mice also displayed a three-fold increase of platelet count, and two-fold increase of platelet volume indicative of macrothrombocytosis (Fig. 2F, G), and a mild leukocytosis driven by an increase of neutrophils (Fig. 2H, Supplementary Table 2). Plasma analysis revealed that Hum mice had elevated iron levels (WT, 44µmol/L; Tg, 49µmol/L; Hum, 107µmol/L; Fig. 2I). This was accompanied by a eight-fold increase in ferritin (Fig. 2J), and modestly elevated transferrin (WT, 1.38g/L; Tg, 1.52g/L; Hum, 1.6g/L; Fig. 2K), suggesting enhanced iron release due to RBC degradation. Bilirubin levels were also doubled in Hum mice compared to WT and Tg littermates (Fig. 2L), further corroborating increased RBC turnover. Bile acid levels, however, remained consistent across genotypes, indicating no hepatic inflammation. Taken together, these data are consistent with accelerated red blood cell destruction rather than impaired iron availability.

**Figure 2.**
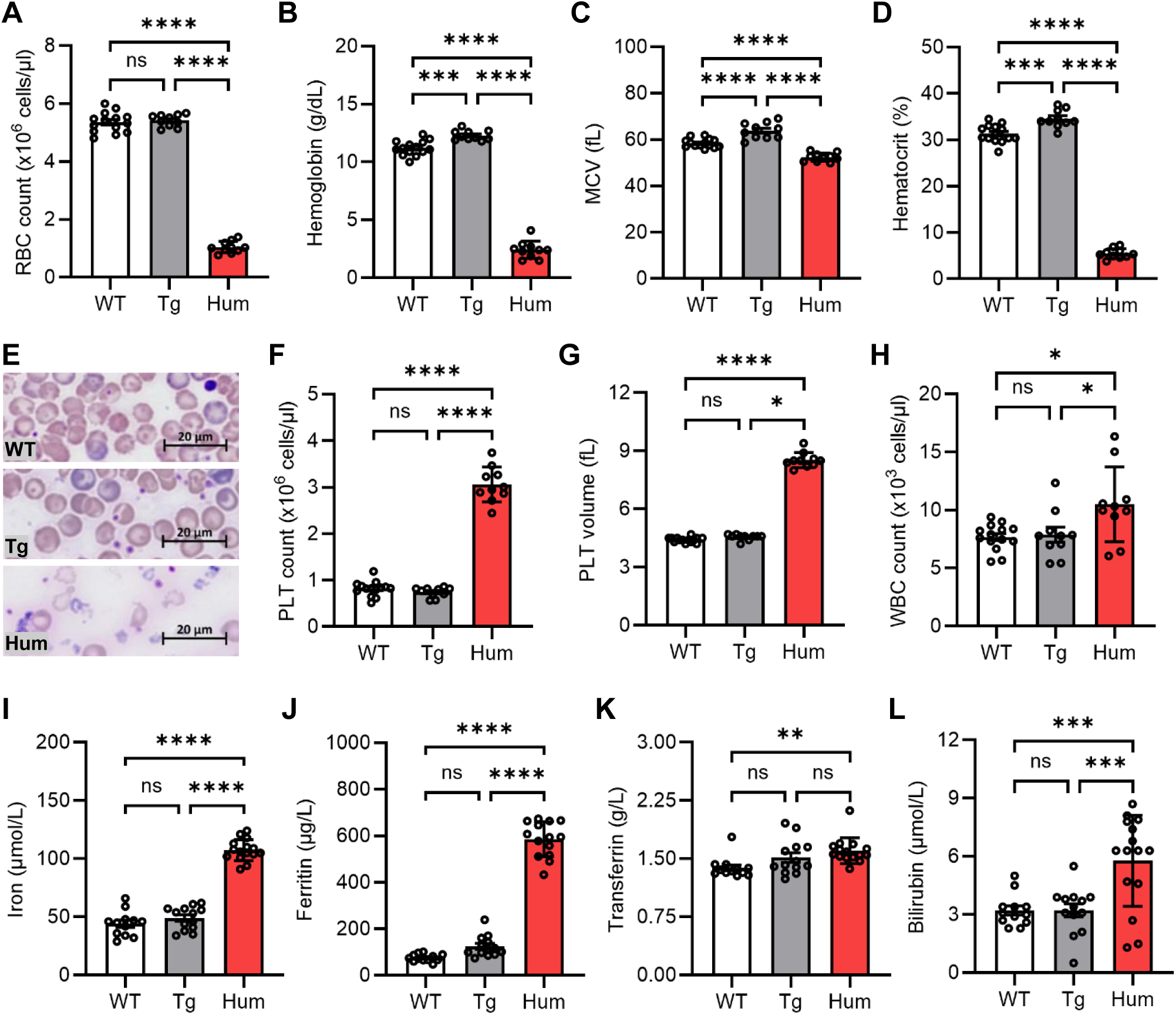
Humanized mice exhibit hemolytic anemia and macrothrombocytosis. (**A**-**H**) Whole blood analysis. RBC: red blood cell; MCV: mean corpuscular volume; PLT: platelet; WBC: white blood cell. (**E**) Representative images of blood smears stained with hematoxylin and eosin. (**I**-**L**) Plasma analysis. Data are mean ± SD, ns: not significant, \**P* < 0.05, \*\**P* < 0.01, \*\*\**P* < 0.001, \*\*\*\**P* < 0.0001. (**A**-**D**, **F**, **H**-**J, L**) One-way ANOVA + Tukey’s post-hoc; (**G**, **K**) Kruskal-Wallis + Dunn’s post-hoc.

### Ultrastructural analysis reveals extensive erythrocyte fragmentation and abnormal platelet morphology

To further investigate the hematological anomalies of Hum animals, we performed scanning electron microscopy (SEM) on PB samples (Fig. 3, Supplementary Table 5). The SEM analysis confirmed our hematological findings, revealing in humanized samples a severe reduction in normal RBC percentage (WT, 76.6%; Tg, 76.3%; Hum, 11.2%; Fig. 3D) and an increase in platelet percentage (WT, 7.5%; Tg, 6.6%; Hum, 16.6%; Fig. 3F). Across all genotypes, 15 to 20% of RBCs displayed abnormal morphology, consisting predominantly of echinocytes in WT and Tg samples, and microcytes and schistocytes in Hum samples (Fig. 3E, H-K). Notably, Hum mice exhibited a high proportion of RBC fragments (51.8%; Fig. 3G), indicative of extensive cell fragmentation (Fig. 3L-M). Finally, we observed that platelets in Hum PB samples were predominantly large, irregularly shaped, and with an elongated morphology (Fig. 3N). These ultrastructural abnormalities provide direct evidence that human DNM2 expression compromises membrane integrity and/or mechanical stability of late erythroid and megakaryocytic lineages.

**Figure 3.**
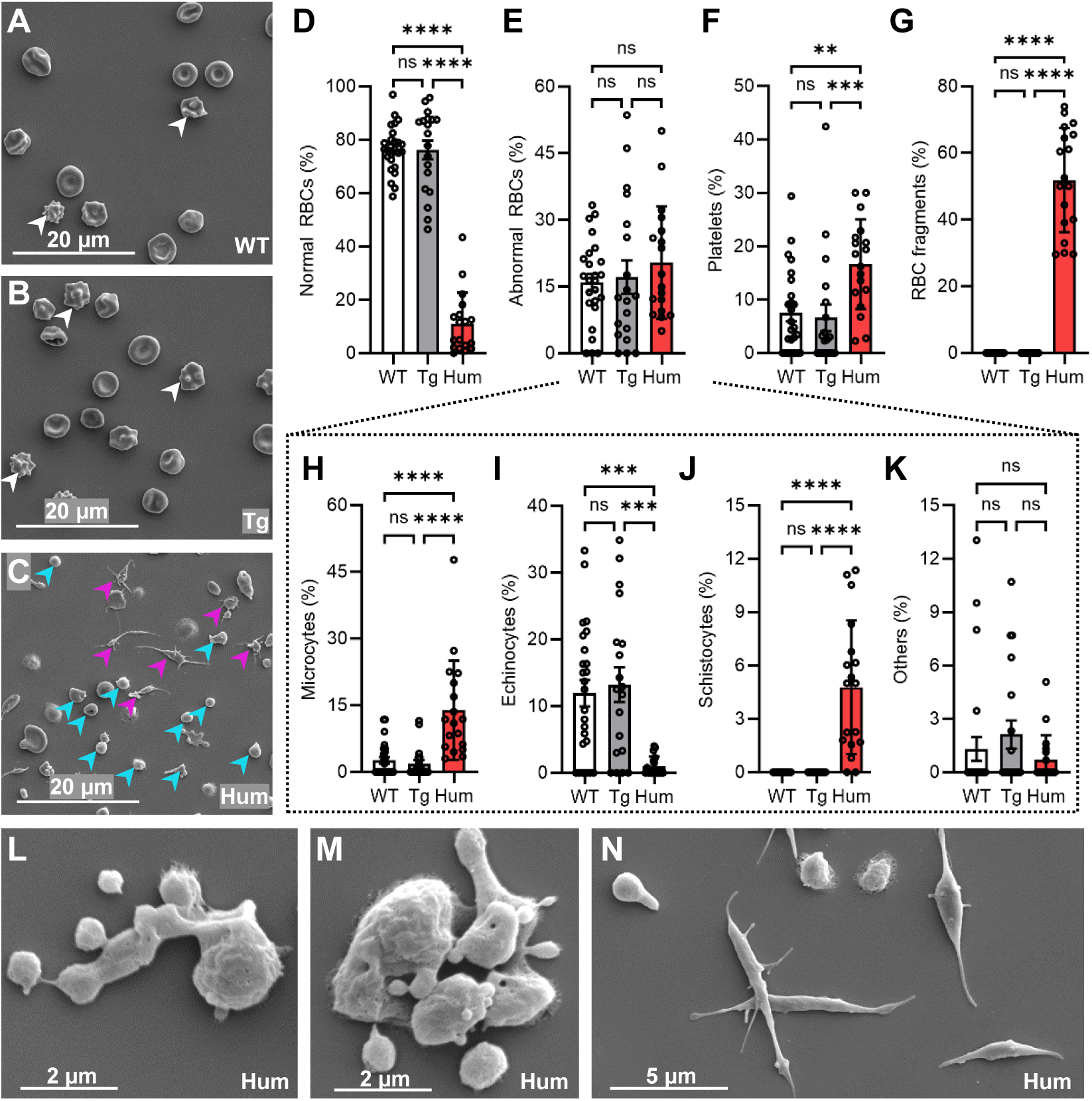
Scanning electron microscopy (SEM) reveals severe abnormalities in RBCs and platelets from *DNM2* humanized mice. (**A**-**C**) SEM representative images of peripheral blood (PB) samples at 2,000x magnification. White arrowheads indicate echinocytes, magenta arrowheads denote platelets, and cyan arrowheads highlight RBC fragments. (**D**-**K**) Quantification of PB cells and fragments (%). (**H**-**K**) Classification of abnormal RBCs (%). (**H**-**I**) Quantification of microcytes, echinocytes, schistocytes (%). (**K**) Quantification of all other poikilocytosis combined including spherocytes, dacrocytes, pyknocytes, and drepanocytes (%). (**L**-**M**) High-magnification SEM images (15,000x) showing RBC fragmentation in Hum samples. (**N**) SEM image at 10,000x illustrating irregular platelet size and morphology in Hum samples. Data are mean ± SD, ns: not significant, \*\**P* < 0.01, \*\*\**P* < 0.001, \*\*\*\**P* < 0.0001. Kruskal-Wallis + Dunn’s post-hoc.

### Fetal liver hematopoiesis is preserved in DNM2-humanized embryos

To determine whether hematopoietic defects arise during embryogenesis, we performed blood smear analysis and RT-qPCR on fetal liver which is the primary site of hematopoiesis between E12.5 and E16.5^35,36^ (Fig. 4, Supplementary Tables 1 & 6). Analysis of blood cell populations at E14.5 revealed comparable distributions across genotypes, characterized by a predominance of mature RBCs (WT, 62.6%; Tg, 57%; Hum 59.7.2%; Fig. 4A-G). Consistent with this, globin expression analysis (Fig. 4H-J) demonstrated a normal fetal-to-adult hemoglobin switch, with adult hemoglobin being the predominant form in all groups (WT, 99%; Tg, 99.5%; Hum 98.9%; Fig. 4K). These findings indicate that human DNM2 supports fetal hematopoiesis, and that hematopoietic failure emerges specifically during postnatal bone marrow–dependent erythropoiesis.

**Figure 4.**
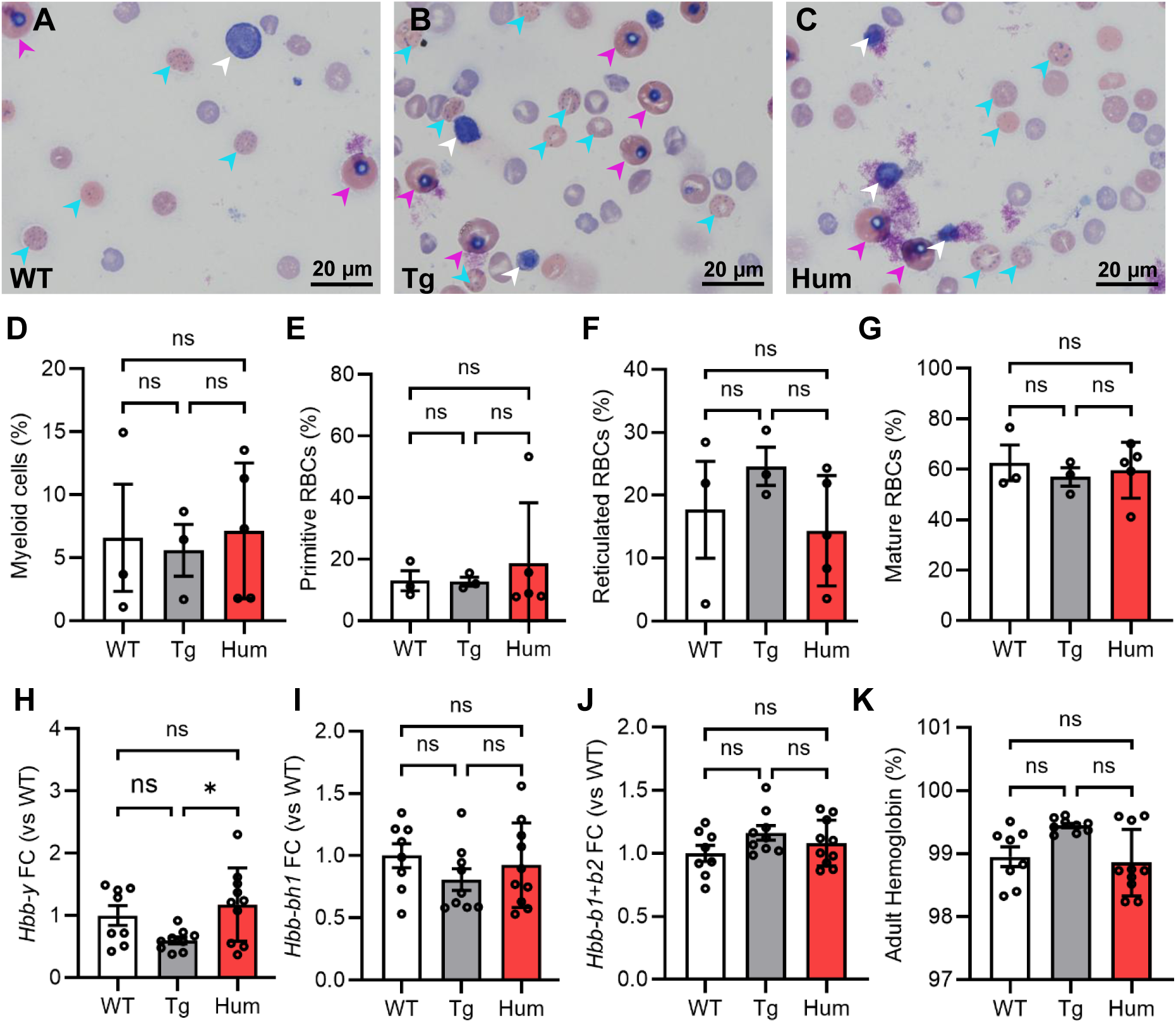
Humanized embryos show normal fetal hematopoiesis at embryonic stage E14.5. (**A**-**C**) Representative images of blood smears after May-Grünwald Giemsa staining. White arrowheads indicate myeloid cells, magenta arrowheads denote primitive RBCs originating from the yolk sac, and blue arrowheads highlight reticulated RBCs. (**D**-**G**) Quantification of cell populations from liver-derived blood smears (% cells). (**H**-**J**) RT-qPCR fold change (FC) results for fetal (*Hbb-y*, *Hbb-bh1*) and adult (*Hbb-b1*, *Hbb-b2*) hemoglobin synthesis–related genes measured in fetal liver cells. (**K**) Adult hemoglobin fraction (%). Data are mean ± SD, ns: not significant, \**P* < 0.05. (**D**, **F**-**J**) One-way ANOVA + Tukey’s post-hoc; (**E**, **K**) Kruskal-Wallis + Dunn’s post-hoc.

### DNM2 humanization causes a block in erythroblast maturation in bone marrow and spleen

To provide a mechanistic insight into the cause of erythrocyte defects, we performed flow cytometry analyses on bone marrow (BM), spleen, and peripheral blood (PB) samples in P12 animals (Fig 5, Supplementary Fig. 5 & Table 7). In the BM, a reduction in cell count was observed in Hum mice (Fig. 5A), which correlates with the decrease in body mass of the animals (Fig. 1F). We first analyzed hematopoietic progenitor and stem cell populations and found that frequencies of multipotent progenitors (Lin^-^Sca1^+^c-Kit^+^; LSK; Fig. 5B) decreased slightly in Hum compared to Tg samples, while no changes in frequencies of common lymphoid progenitors (CLP; Fig. 5C), common myeloid progenitors (CMP; Fig. 5D), granulocyte-monocyte progenitors (GMP; Fig. 5E), and megakaryocytic-erythroid progenitors (MEP; Fig 5F). We then used the erythroblast markers CD71 and Ter119 to distinguish erythroid populations according to their maturation stage (Fig. 5G): proerythroblast (ProE, CD71^+^ Ter119^-^), basophilic erythroblast (EryA, CD71^+^ Ter119^+^ high FSC), polychromatophilic erythroblast (EryB, CD71^+^ Ter119^+^ low FSC), and orthochromatic erythroblast (EryC, CD71^-^ Ter119^+^ low FSC). We observed a tenfold increase of ProE cells in Hum animals (WT, 6.8%; Tg, 5.6%; Hum 69.5%; Fig. 5H), while other erythroblast populations were decreased in Hum samples compared to the other genotypes (Fig. 5I-K). While WT and Tg animals predominantly displayed EryB and EryC populations, the EryC subset was nearly absent in Hum mice (WT, 20%; Tg, 29.5%; Hum 1.89%; Fig. 5K).

**Figure 5.**
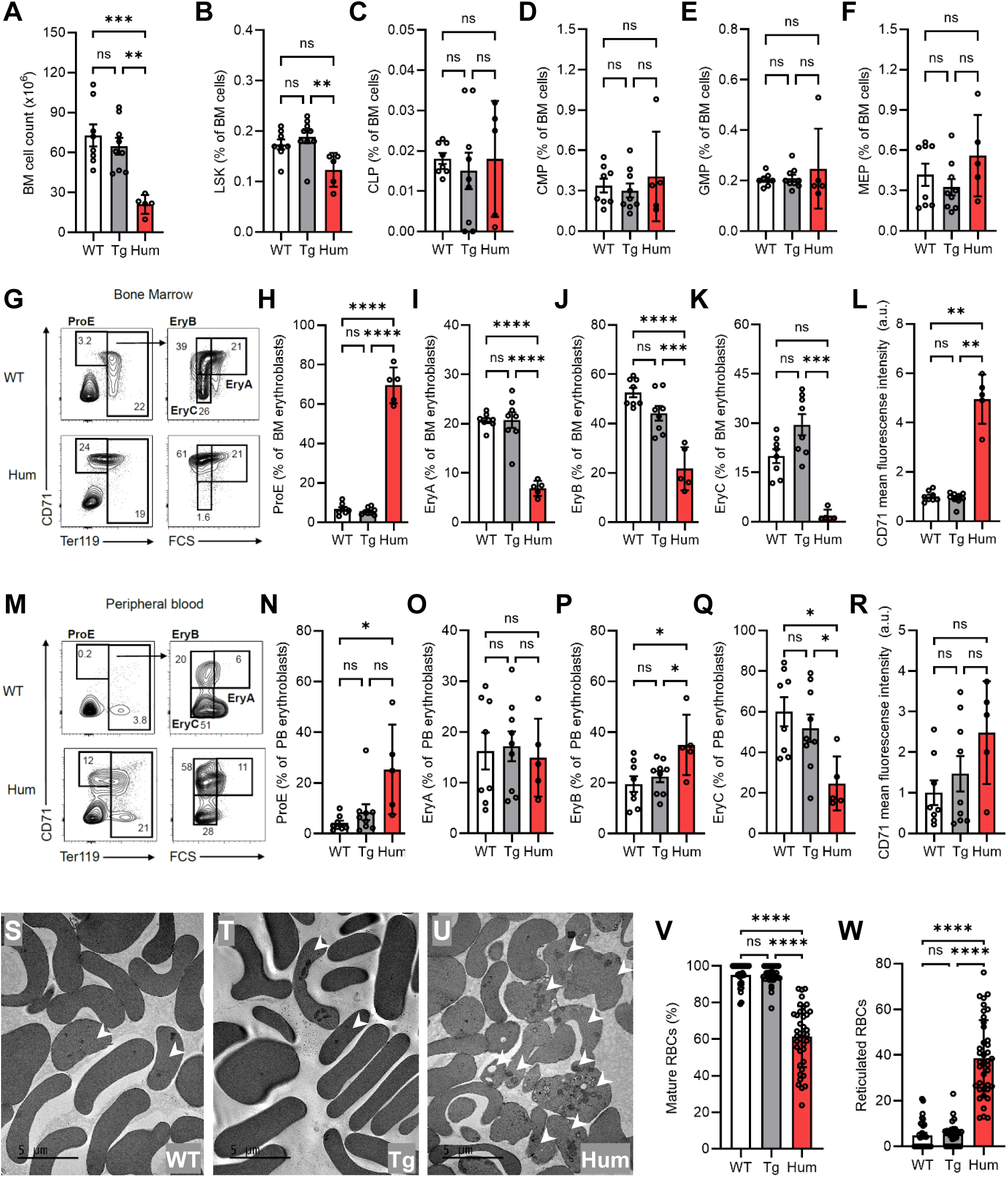
Impaired erythroblast maturation in the bone marrow (BM) and peripheral blood (PB) of humanized mice. (**A**-**L**) BM analysis. (**M**-**W**) PB analysis. (**A**) Cell count derived from BM samples (x10^6^ cells). (**B**-**F**) LSK/LK hematopoietic progenitors (% cells). LSK: Lin^-^Sca1^+^c-Kit^+^ hematopoietic stem and progenitor cells. CLP: Common lymphoid progenitors. CMP: Common myeloid progenitors. GMP: Granulo-monocyte progenitors. MEP: Megakaryo-erythrocyte progenitors. (**G**, **M**) Representative flow cytometry plots using CD71 and Ter119 expressions with the indicative percentage of cells. (**H**-**K**, **N**-**Q**) Erythroblast populations (% erythroblasts). ProE: proerythroblasts. EryA: basophilic erythroblasts. EryB: polychromatophilic erythroblasts. EryC: orthochromatic erythroblasts. (**L**, **R**) CD71 mean fluorescence intensity in erythroblasts (arbitrary units). (**S**-**U**) Transmission electron microscopy (TEM) representative images of PB samples at 1,500x magnification with white arrowheads showing reticulocytes. (**V**-**W**) Quantification of PB mature and reticulated RBCs (% cells). Data are mean ± SD, ns: not significant, \**P* < 0.05, \*\**P* < 0.01, \*\*\**P* < 0.001, \*\*\*\**P* < 0.0001. (**A**, **C**, **H**-**J, O**-**R**) One-way ANOVA + Tukey’s post-hoc; (**B**, **D**, **E**, **F**, **K**-**L**, **N**, **V**-**W**) Kruskal-Wallis + Dunn’s post-hoc.

A comparable distribution of erythroblast subpopulations was observed in the humanized spleen (Supplementary Fig. 5), indicating that erythropoiesis was impaired in both medullary and extramedullary sites. Notably, although spleen masses were similar between WT and Hum mice (Fig. 1G), the cell number obtained from total spleens was markedly reduced in the humanized group (WT, 182.10^6^; Tg, 186.10^6^; Hum 37.10^6^; Supplementary Fig. 5A). These findings corroborate our histological data and suggest that the splenomegaly in Hum animals likely results from defective RBC recycling. Together, these findings identify a specific block in late erythroid maturation rather than a global hematopoietic defect.

### Humanized erythroblasts display increased CD71 surface expression and delayed maturation in peripheral blood

To confirm the defect in erythroid maturation and assess the role of DNM2 in this process, we focused on transferrin receptor internalization, and studied PB cells. Importantly, cell surface expression of the transferrin receptor CD71 in BM cells was fivefold higher in humanized erythroblasts relative to WT and Tg (WT, 1; Tg, 0.94; Hum, 4.95; arbitrary unit; Fig. 5L). While *Tfrc* expression remained stable across fetal liver samples, it was twofold higher in postnatal Hum BM relative to WT and Tg mice (WT, 1; Tg, 0.98; Hum, 2.21; arb. unit; Supplementary Fig. 6A). Comparable CD71 cell surface expression (Supplementary Fig. 5L) and transcriptional upregulation of its corresponding *Tfrc* gene (Supplementary Fig. 6B) were observed in the humanized spleen. These findings are consistent with impaired transferrin receptor turnover in postnatal erythroblasts expressing human DNM2, in line with the established role of dynamin 2 in clathrin-mediated endocytosis of the transferrin receptor^37^.

The erythroid maturation defect extended to the PB samples (Fig. 5M-Q). The relative frequencies of EryA and EryB were comparable among genotypes; however, Hum animals exhibited again an increased proportion of proE (WT, 4.15%; Tg, 8.48%; Hum 25.4%; Fig. 5N). Consequently, the mature EryC cell population, which predominated in WT and Tg mice, was substantially reduced in Hum animals (WT, 60%; Tg, 51.9%; Hum 24.7%; Fig. 5Q). To further discriminate mature erythrocytes from reticulocytes, which retain residual organelles, we performed transmission electron microscopy (TEM) on PB samples (Fig. 5S-W & Supplementary Table 8). In agreement with the flow cytometry data, PB from WT and Tg mice consisted predominantly of mature erythrocytes (WT, 95.1%; Tg, 94.5%; Hum 61.4%), whereas PB from Hum mice contained a higher proportion of reticulocytes, representing over one-third of the total erythroid population.

To assess whether the localization of the human DNM2 protein differs from its endogenous counterpart and could underlie the cytometric observations, we conducted immunofluorescence staining on BM cytospin preparations (Supplementary Fig. 7). Our results revealed similar localization of both proteins in nucleated hematopoietic cells, indicating 1) no mislocalization of the human protein and 2) strong reduction of the expression of DNM2 protein in enucleated erythrocytes. Thus, the phenotype arises from functional impairment during erythroblast stages, not from ectopic DNM2 persistence in erythrocytes.

Collectively, these observations support a model in which expression of human DNM2 impairs late erythroid differentiation, likely through disruption of membrane trafficking processes required for transferrin receptor turnover, thereby impairing terminal erythroid maturation.

### Species-specific divergence of the DNM2 proline-rich domain rewires the hematopoietic interactome

We next investigated the molecular basis for the species-specific differences between human and mouse DNM2 that may contribute to the erythroid phenotype and anemia in humanized mice. Since most of the amino acid (aa) differences between human and mouse DNM2 proteins are located in the proline-rich domain (PRD; Fig. 6A) spanning residues P747 to D870, we decided to investigate the interactomes of PRDs from both species using native holdup experiments. However, due to the high hydrophobicity of the domain, full-length synthesis of PRDs was not feasible. Instead, we synthesized two peptides per species: Q758 to A783 and A820 to I842 (similar numbering in mouse and human), each peptide encompassing 4 substitutions, thereby covering 8 of the 12 aa changes between human and mouse PRDs (Fig. 7B). The biotinylated peptides were used as a bait on streptavidin resin which was incubated with cell lysate from a mouse hematopoietic progenitor cell line (HPC-7). High DNM2 expression in the HPC-7 cell line was confirmed both by Western blot analysis (Supplementary Figure 8), and the HPC-7 proteome (Supplementary Table 9). Mass spectrometry analysis of unbound fractions from HPC-7 cell extracts was performed to determine protein binding affinities. A total of 8,925 proteins were identified across all samples (Supplementary Tables 10-11), and after filtering, 5,883 proteins were quantified in this analysis (Supplementary Tables 12-15). While mouse peptides M1 and M2 interacted with RNF11, IFT25, PLAC8, and GRB2 proteins, human peptides H1 and H2 bound to PP6R2, ING2, GRB2, UBS3A (encoded by the *Ubash3a* gene), MAD3 (encoded by the *Mxd3* gene), TMUB1, and IPKG proteins (Fig. 6C & Supplementary Table 16). Notably, GRB2, a SH3 domain-containing protein implicated in signal transduction and a known interactor of DNM2^38^, was the only shared interactor between the mouse and human peptides (H2 & M2) whereas RNF11 interacted with both M1 and M2 mouse peptides.

**Figure 6.**
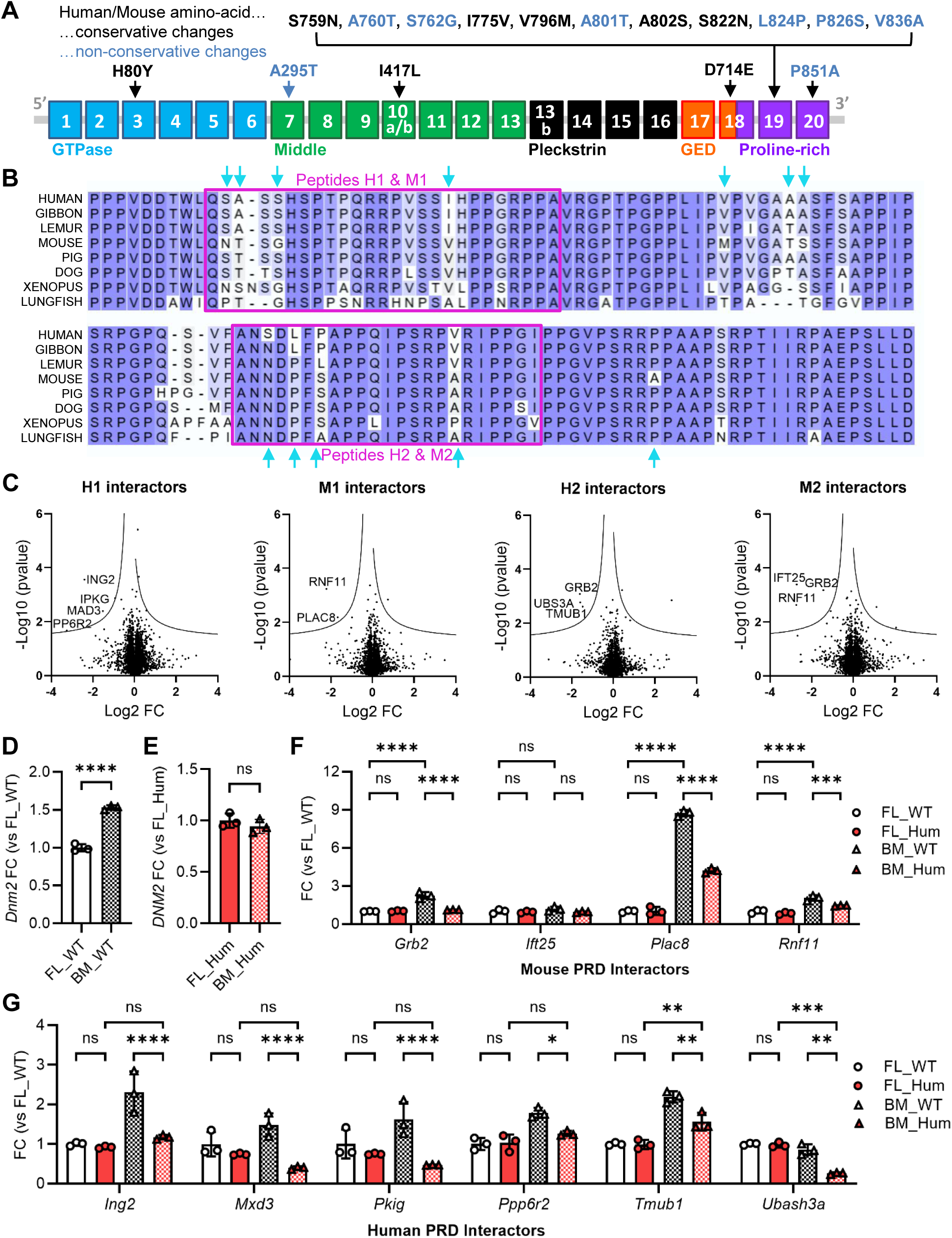
Identification of human- and mouse-specific interactors of the DNM2 Proline-Rich Domain (PRD) using peptide-based native holdup (nHU) assay. (**A**) DNM2 protein domains and amino acid (aa) discrepancies between humans and mice. Of the 870 aa, only 16 differ including 12 aa located in the proline rich-domain. (**B**) Sequence alignment of DNM2 PRD domains across tetrapod species obtained from Uniprot. Magenta squares highlight peptide sequences used in the holdup assay and cyan arrows indicate amino acid variations between human and mouse species. (**C**) Volcano plot of the nHU-MS experiment performed with human and mouse peptides on total HPC-7 extract. Identified interaction partners are indicated by their protein name on the graph. (**D**-**G**) RNA fold change (FC) measurements of *Dnm2*, *DNM2*, and genes coding for DNM2-PRD interactors in E14.5 fetal liver (FL) and P12 bone marrow (BM) samples in arbitrary units (FL_WT mean =1), Shown in **F**, *Grb2* was not included in **G**. Data are mean ± SD, \**P* < 0.05, \*\**P* < 0.01, \*\*\**P* < 0.001, \*\*\*\**P* < 0.0001. (**D**, **E**) Student’s *t*-test. (**F**, **G**) Two-way ANOVA + Tukey’s post-hoc.

**Figure 7.**
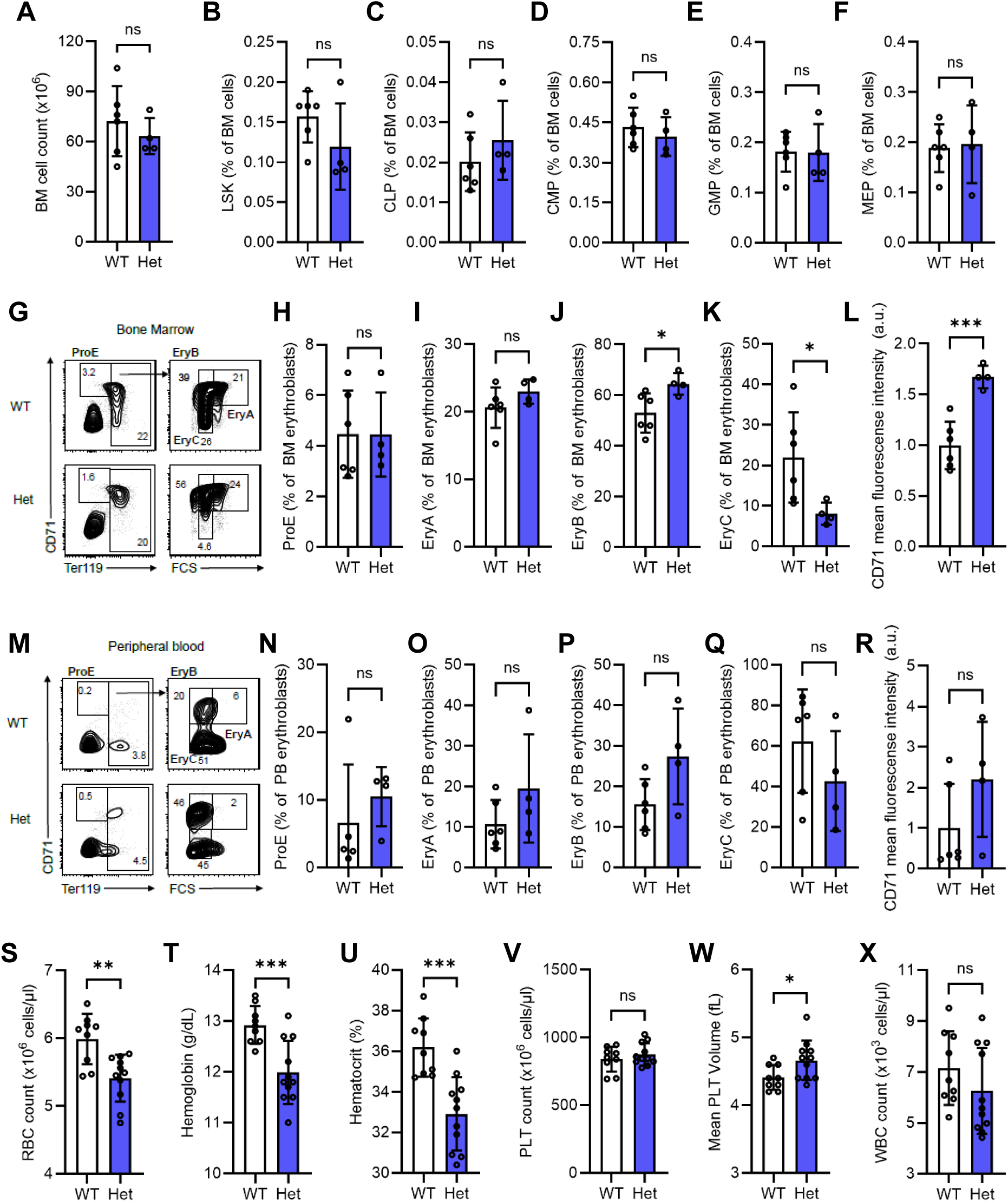
*Dnm2* haploinsufficiency results in reduced EryC populations and elevated CD71 surface expression in the bone marrow (BM), with concomitant decreases in erythrocyte counts and hemoglobin levels in peripheral blood (PB). (**A**-**R**) Flow cytometry analysis. (**A**-**L**) BM analysis. (**M**-**R**) PB analysis. (**A**) Cell count derived from BM samples (x10^6^ cells). (**B**-**F**) LSK/LK hematopoietic progenitors (% cells). (**G**, **M**) Representative flow cytometry plots using CD71 and Ter119 expressions with the indicative percentage of cells. (**H**-**K**, **N**-**Q**) Erythroblast populations (% erythroblasts). (**L**, **R**) CD71 mean fluorescence intensity in erythroblasts (arbitrary units). (**S**-**X**) Whole blood analysis. RBC: red blood cell; MCV: mean corpuscular volume; PLT: platelet; WBC: white blood cell. Data are mean ± SD, ns: not significant, \**P* < 0.05, \*\*\**P* < 0.001. (**A**, **C**-**F**, **H**-**J**, **L**, **O**-**Q**, **S**-**X**) Student’s *t*-test; (**K**) Welch’s *t*-test; (**B**, **N**, **R**) Mann-Whithey test.

Given the normal hematopoiesis in the fetal liver (FL) at E14.5 and its impairment in BM at P12 in Hum animals, we compared the expression of *DNM2* human and mouse genes and their PRD interactors in both tissues between WT and Hum mice. We first observed that *Dnm2* endogenous expression is higher in the BM than in the FL and that *DNM2* transgenes are highly stable and consistently expressed (Fig. 6D, E). While gene expression levels were comparable between WT and Hum FL samples, all genes except *Ift25* exhibited reduced expression in the Hum BM samples relative to WT BM samples (Fig. 6F, G). Among the genes encoding mouse peptide interactors, *Grb2*, *Rnf11*, and *Plac8* expressions were higher in BM than in FL, with an eightfold increase for the latter (WT FL, 1; Hum FL, 1; WT BM, 8.7; Hum BM, 4.2; arb. unit). Among genes encoding human peptide interactors, only *Tmub1* displayed higher expression in BM compared to FL whereas *Ubash3a* showed lower expression.

Together, these data indicate that evolutionary divergence of the DNM2 PRD reshapes interaction networks critical for postnatal erythropoiesis, ultimately leading to severe hemolytic anemia in humanized mice.

### *Dnm2* haploinsufficiency phenocopies erythroid and erythrocyte defects with reduced severity

We observed both gain and loss of specific interactors comparing human and mouse DNM2 PRD. To determine whether the phenotypes observed in *DNM2*-humanized animals reflected a gain- or loss-of-function effect of human DNM2 compared to mouse DNM2, we performed flow cytometry experiments on samples from *Dnm2* heterozygous (Het) mice mimicking a partial DNM2 loss-of-function (Fig. 7, Supplementary Fig. 9 & Table 17). In the BM, hematopoietic progenitor and stem cell populations were comparable among groups (Fig. 7B-F), while we observed an increase of EryB and a decrease of EryC percentages in Het samples compared to WT (Fig. 7G-K). Notably, cell surface expression of the transferrin receptor CD71 was higher in Het mice compared with controls, both on BM erythroblasts (WT, 1; Het, 1.67; arb. unit; Fig. 7L) and spleen erythroblasts (WT, 1; Het, 1.35; arb. unit; Supplementary Fig. 9G). This increase was less pronounced than that observed in Hum animals and was accompanied by *Tfrc* transcriptional upregulation in the spleen (WT, 1.57; Het, 1.86; arb. unit; Supplementary Fig. 6B), but not in the BM. DNM2 haploinsufficiency had no effect on the overall erythroblast composition or CD71 surface expression in the PB (Fig. 7M-R). However, whole blood analysis revealed that *Dnm2*^+/-^ mice show reduced RBC count and hemoglobin levels, along with increased platelet volume (Fig. 7S-X; Supplementary Fig. 18). These results indicate that *Dnm2* heterozygous mice exhibit an intermediate phenotype relative to *DNM2*-humanized animals, supporting a loss-of-function mechanism and consistent with impaired PRD-dependent interaction networks of the human protein in this murine context.

## DISCUSSION

In this study, to better understand the physiological role of human DNM2, we created and characterized a humanized mouse model, and show that complete replacement of murine DNM2 by human DNM2 supports embryonic development but fails to sustain postnatal hematopoiesis, leading to severe hemolytic anemia and early lethality. We identify a lineage-specific block in late erythroid maturation characterized by accumulation of immature erythroblasts, reticulocytosis, red blood cell fragmentation, and compensatory extramedullary hematopoiesis, despite preserved hematopoietic progenitors and normal fetal liver erythropoiesis. This defect is associated with elevated CD71 transferrin receptor surface expression, consistent with impaired membrane trafficking during terminal erythropoiesis. At the molecular level, we uncover extensive species-specific divergence of the DNM2 proline-rich domain, resulting in rewired interaction networks in hematopoietic progenitors. Finally, analysis of *Dnm2* haploinsufficient mice reveals a graded, intermediate phenotype, supporting a dosage-sensitive loss-of-function mechanism rather than a toxic gain-of-function effect of human DNM2 in the murine hematopoietic context.

### Human DNM2 rescues embryonic lethality through conserved core functions and the GRB2 axis

A central finding of this study is that complete replacement of murine *Dnm2* by human *DNM2* rescues embryonic development despite leading to postnatal hematopoietic failure. Whereas constitutive deletion of *Dnm2* causes early embryonic lethality^28,29^, human DNM2 preserves essential dynamin functions sufficient to sustain embryogenesis, indicating strong conservation of core molecular activities. Our interactome analyses provide a mechanistic basis for this paradox by identifying GRB2 as the only proline-rich domain interactor conserved between human and mouse DNM2. GRB2 is a critical adaptor linking receptor tyrosine kinase signaling to Ras–MAPK activation^39,40^ and has been shown to associate with DNM2, coupling clathrin-mediated endocytosis to growth factor–dependent signal transduction^38^. Consistent with this, both *Dnm2* and *Grb2* knockout mice exhibit early embryonic lethality^28,29,41^, underscoring the importance of this axis during development. We propose that preservation of the GRB2–DNM2 interaction constitutes a minimal, evolutionarily conserved module sufficient for embryonic viability. In contrast, postnatal tissues—particularly the hematopoietic system—require expanded and lineage-specific PRD-dependent interaction networks that are not conserved between species, thereby revealing a functional stratification within a ubiquitously essential protein.

### DNM2 is essential for postnatal erythropoiesis through regulation of transferrin receptor turnover

Our findings identify DNM2 as a critical regulator of postnatal erythropoiesis and reveal a previously unrecognized requirement for species-specific DNM2 function during late erythroid maturation. In DNM2-humanized mice, erythropoiesis is profoundly disrupted after birth, with a marked delay in the maturation of proerythroblasts into erythroblasts in both the bone marrow and spleen, reticulocytosis, red blood cell fragmentation, and compensatory extramedullary hematopoiesis, accompanied by macrothrombocytosis. In addition, plasma analysis demonstrated significant elevations in iron, ferritin, and bilirubin levels in Hum samples, indicative of red blood cell (RBC) degradation and heightened hemolytic activity. These defects occur despite preserved hematopoietic stem and progenitor compartments and normal fetal liver erythropoiesis, indicating a lineage- and stage-specific failure that emerges with bone marrow–dependent erythropoiesis.

A central feature of this phenotype is the strong accumulation of the transferrin receptor CD71 at the erythroblast surface, together with transcriptional upregulation of the corresponding *Tfrc* gene, consistent with impaired transferrin receptor turnover. This finding is mechanistically aligned with the established role of dynamin 2 in clathrin-mediated endocytosis^42^ and highlights the exceptional sensitivity of erythroblasts to perturbations in membrane trafficking, given their high iron flux and extensive membrane remodeling during differentiation^43^.

To date, no RBC disorder has been reported in patients with *DNM2* mutations; however, mouse model studies have highlighted the role of DNM2 in hematopoiesis. In an N-ethyl-N-nitrosourea–induced mouse model, a heterozygous V235G mutation within the GTPase domain of *DNM2* impaired its enzymatic activity and led to mild microcytic anemia, despite RBC counts and serum levels of iron, ferritin, and transferrin remained within normal ranges^44^. Our humanized model extends these observations by demonstrating that, while human DNM2 is sufficient to sustain embryonic endocytic requirements, it is inadequate to support the specialized trafficking demands of postnatal erythropoiesis. Together, these results position DNM2 as a key node in erythroid membrane trafficking whose precise functional tuning is essential for terminal erythroid maturation and red blood cell stability.

### PRD divergence rewires interaction networks and drives a dosage-sensitive loss-of-function phenotype

Although human and mouse DNM2 share 98% sequence identity, comparative sequence analysis reveals that the majority of amino acid differences are concentrated within the C-terminal proline-rich domain (PRD), a low-complexity region that mediates interactions with SH3 domain–containing proteins^7^ and appears to function as a rapidly evolving regulatory interface. Indeed, of the 16 amino acid substitutions distinguishing human and murine DNM2, 12 localize to the PRD, including most non-conservative changes, whereas the catalytic GTPase and membrane-binding domains remain highly conserved. The unique substitution located in the GTPase domain, His80Tyr, is frequently reported in the gnomAD database for human genomic variants, with 3,000 heterozygous and a single homozygous carrier (Supplementary Table 19). This asymmetric divergence suggests that evolutionary variation preferentially targets regulatory interaction surfaces rather than enzymatic core functions. Consistent with this notion, native holdup proteomic analyses revealed an extensive rewiring of PRD-dependent interaction networks in hematopoietic progenitors (HPC-7), with GRB2 emerging as the sole conserved interactor between species. In contrast, several murine-specific PRD partners, namely IFT25, RNF11 and PLAC8, were lost in the humanized context, while human PRD peptides gained interactions with proteins such as UBS3A, TMUB1, and MAD3, which are implicated in immune signaling and transcriptional regulation. Comparison of fetal liver and postnatal bone marrow expression profiles highlighted PLAC8 as a particularly compelling candidate, as it is strongly enriched in postnatal bone marrow, associated with myeloid differentiation^45^ and immune activation^46,47^, and selectively interacts with the murine—but not human—DNM2 PRD. Loss of this interaction may therefore contribute to the erythroid-specific phenotype observed in humanized mice. However, the phenotype observed in DNM2-humanized mice is more likely to result from the combined loss and reorganization of multiple PRD-dependent interactions, rather than from disruption of a single interaction.

Genetic dosage experiments further resolve the functional consequences of this interactome rewiring: *Dnm2* heterozygous mice display intermediate erythroid defects, with increased CD71 surface expression and mildly impaired erythroblast maturation in the BM, along with reduced RBC count and hemoglobin level in the PB. These findings support a dosage-sensitive loss-of-function mechanism rather than a toxic gain-of-function effect of human DNM2.

Lastly, comparative sequence analysis across tetrapod species reveals notable amino acid differences within the PRD of DNM2 (Fig. 6B). The PRD displays a distinct pattern of evolutionary divergence in great apes, consistent with primate-specific adaptation of a regulatory interaction domain with potential tissue-specific consequences.

### Conclusions

Together, our findings demonstrate that evolutionary divergence of DNM2 has lineage-specific functional consequences, revealing how subtle variation in regulatory domains can profoundly alter physiological outcomes. We show that divergence of the DNM2 proline-rich domain rewires interaction networks required for postnatal erythropoiesis, despite preservation of core functions sufficient for embryonic development. This identifies PRDs as rapidly evolving context-dependent adaptors that tune ubiquitous proteins to tissue-specific demands. More broadly, our work highlights how evolutionary rewiring of protein interaction interfaces can expose hidden vulnerabilities in humanized models and underscores the need to consider regulatory domain compatibility—not only core function conservation—when designing gene replacement strategies.

## MATERIALS AND METHODS

### Sex as a biological variable

Our study examined male mice because the human DNM2 transgenes were integrated on the Y chromosome.

### Mouse lines, genotyping, and ethical statement

The creation and characterization of *Dnm2+/-* animals were described previously^29^. Briefly, we introduced LoxP sites flanking exon 8 and crossed floxed mice with CMV-Cre mice to generate Dnm2+/- animals. Deletion of exon 8 is predicted to induce a frameshift and premature stop codon in exon 9. The DNM2 transgenic (Tg) mouse line was generated at the ICS (Institut Clinique de la Souris; https://ics-mci.fr/en/). The bacterial artificial chromosome (BAC) RP11-20N24 (Supplementary Fig. 1A) was ordered from CHORI BAC. This BAC bares the human chromosome 19 genomic region spanning from 10,790,427 to 10,952,432 (GRCh37) which includes human DNM2 gene (ENSG00000079805) and passenger genes (QTRT1 ENSG00000213339 and TMED1 ENSG00000099203). The regions containing exons were amplified by PCR and sequenced by Sanger sequencing. This confirmed the presence of the entire gene on the BAC. The BAC was microinjected into C57BL/6N fertilized eggs and then injected eggs were implanted into the oviducts of pseudopregnant foster mice. One founder was breed for germ line transmission and one line was established. Droplet digital PCR copy counting confirmed the presence of 2 full copies of the BAC (Supplementary Fig. 1B, C). The transgenes (Tg) are located on the Y chromosome as only males were found positive. Tg*DNM2* animals were crossed with *Dnm2^+/-^*to generate *Dnm2^+/-^*Tg*DNM2* which were crossed with *Dnm2^+/-^*to generate *Dnm2^-/-^*Tg*DNM2* (Hum) and *Dnm2^+/+^*Tg*DNM2* (Tg) littermates (Supplementary Fig. 2A). Wild-type (WT) animals were generated separately by mating *Dnm2^+/-^*animals together. *Dnm2* WT and exon-8 deletion alleles were identified by using 5′-TGT CGC TCC TGT GGG ACC GAG G-3′ (forward) and 5′-TCC TGC CTC AAC TCA CAC AAC TCT GC-3′ (reverse) primer pair. Polymerase chain reactions (PCRs) amplified specific products from deletion and WT alleles of 553 and 1517 bp, respectively. DNM2 transgene genotyping was done using 5′-GTC CAT CTG GTC CCA GCT TTC-3′ (forward) and 5′-CAC CCT GGA CAG CTA GCC AG-3′ (reverse) primer pair. PCRs amplified specific transgene product of 553 bp. Phire Tissue Direct PCR Master Mix (Thermo Scientific, F-170L, MA, US) was used to perform PCRs according to the manufacturer’s instruction with the following program: 98°C/5 min; 35x (98°C/15s, 61°C/15s, 72°C/30s); 72°C/5min. Animals analyzed had a 50% 129S2/SvPas 50% C57BL/6N mixed background.

### Blood collection, and counting

Following sacrifice, peripheral blood (PB) was collected from P12 mice for subsequent hematology, plasma, and biochemistry analyses. PB was collected in EDTA-coated Microvette 500 K3E tubes (Sarstedt, Nümbrecht, Germany) for whole blood analysis and in Li-heparin-coated tubes (Sarstedt) for plasma analysis. A complete blood count was performed on whole blood on an HT5 analyzer (Scil, Altorf, France). Blood chemistry was performed on an OLYMPUS AU-480 automated laboratory work station (Beckmann Coulter, CA, US) with adapted kits and controls. Whole blood analyses were performed on 14 WT, 10 Tg, and 10 Hum samples, while plasma analyses included 12 WT, 13 Tg, and 15 Hum samples. To study *Dnm2* heterozygous mice, whole blood analyses were performed on 9 WT, and 11 *Dnm2*^+/-^ samples.

### Blood smears

PB was collected using hematocrit capillaries, and a few microliters were deposited onto microscope slides. A second slide was used to spread the blood evenly along the entire length of the experimental slide. The smears were air-dried overnight and imaged the following day using a AxioObserver Z1 time-lapse microscope (Zeiss, Oberkochen, Germany). Brightfield images were acquired with a Plan-Apochromat 63×/1.4 oil immersion objective and a Zeiss Axiocam 305 12-bit color camera (5 MP, pixel size: 3.45 µm). For fetal blood quantification, ten images per sample were acquired from 3 WT, 3 Tg, and 5 Hum samples.

### Histology

After PB collection, spleens, livers, gastrocnemius, quadriceps, and sciatic nerves were collected for histology analysis. Spleens, and livers were fixed by immersion in 4% paraformaldehyde (PFA) for 48 h and embedded in paraffin. Longitudinal sections were cut at 5µm, stained with hematoxylin and eosin, and imaged using the NanoZoomer 2HT slide-scanner (Hamamatsu, Japan). Sciatic nerves and gastrocnemius were fixed by immersion in 2.5% glutaraldehyde and 2.5% PFA in cacodylate buffer (0.1 M, pH 7.4), washed in cacodylate buffer for further 30 minutes. The samples were post-fixed in 1% osmium tetroxide in 0.1M cacodylate buffer for 1 hour at 4°C and dehydrated through graded alcohol (50, 70, 90, and 100%) and propylene oxide for 30 minutes each. The tissues were embedded in Epon 812 imbedding resin. Semithin sections were cut at 2µm, contrasted with toluidine blue, and imaged with a Zeiss AxioObserver Z1 microscope. Ultrathin sections of sciatic nerve were cut at 70nm (Leica Ultracut UCT, Wetzlar, Germany), contrasted with uranyl acetate and lead citrate, and examined at 70kv with a Morgagni 268D electron microscope (FEI Electron Optics, Eindhoven, the Netherlands). Images were captured digitally by Mega View III camera (Soft Imaging System, Münster, Germany).

### RNA extraction, cDNA synthesis, and quantitative PCR

Liquid nitrogen snap-frozen quadriceps, fetal livers (FL), and cell pellets from bone marrow (BM) and spleen were mechanically lysed with a Precellys homogenizer (Bertin technologies, Montigny-le-Bretonneux, France) in Tri Reagent (Molecular Research Center TR118, OH, US) and processed according to the manufacturer’s instructions to obtain total RNA. Concentrations were determined by spectrophotometry (Nanodrop 2000, Thermo Fisher Scientific, MA, US). cDNA was produced from RNA using SuperScript IV Reverse Transcriptase (Thermo Fisher Scientific #18090010) according to the manufacturer’s instruction. The list of primers used are listed in Supplementary Table 20. Reverse transcription quantitative PCRs (RT-qPCRs) were performed in a LightCycler 480 (Roche, Basel, Switzerland) in triplicate reactions set up with SYBR Green Master Mix I (Roche, #04707516001, Basel, Switzerland). Gene expression levels were normalized to *Rpl27* for quadriceps samples and *Psmd1* for FL, BM, and spleen samples, using the 2^(-ΔΔCt)^ method. Results are presented as fold change (FC) relative to the mean expression of WT or Hum groups. The adult hemoglobin fraction was calculated as Q_adult_/(Q_adult_+Q_fetal_)*100 with Q=2^(−ΔCt)^, Q_fetal_ = Q_Hbb-y_+Q_Hbb-bh1_ and Q_adult_ = Q_Hbb-b1_+Q_Hbb-b2_. Analyses were performed using quadriceps samples from 9 WT, 9Tg, 8Hum mice; FL samples from 8 WT, 9Tg, 10Hum mice; and 3 BM and spleen samples per genotype. To study *Dnm2* heterozygous mice, 3WT, and 3*Dnm2*^+/-^ BM and spleen samples were used.

### Protein extraction and Western blotting

Liquid nitrogen snap-frozen tissues were mechanically lysed with a Precellys homogenizer in RIPA buffer supplemented with EDTA-free Protease Inhibitor Cocktail (Roche, #11873580001, Basel, Switzerland). 20 mg of proteins per sample was applied to SDS-PAGE gels, and proteins were transferred to nitrocellulose membranes using Trans-blot semi-dry transfer system (Bio-Rad, Hercules, CA, USA). Protein load control was obtained as quantification of Ponceau S staining. The membranes were blocked in 5% milk and incubated overnight with rabbit antibodies directed against human DNM2 (1:1,000, Thermo Fisher Scientific, #PA5-19800), mouse DNM2 (1:10,000, homemade 2865), or GAPDH (1:1000, Sigma-Aldrich, MO, USA, #MAB374) and for 1 h with 1:10,000 secondary antibodies conjugated with horseradish. The membranes were exposed to ECL and scanned in an Amersham Imager 600 (GE Healthcare Life Sciences, MA, US). Blots and total protein (Ponceau S) densities were quantified in ImageJ. The analyses were performed using 7 quadricep samples per genotype for quantification and 1 sample per genotype for other tissues. All unedited gels used for protein quantification and their loading controls are provided in supplementary materials.

### Flow cytometry analysis

BM cells were isolated from femurs, tibias, pelvic bones, and sternum. PB samples were treated with 1% dextran for 30 min at 37°C to allow erythrocyte sedimentation, and the upper cell suspension was collected for staining. Cells were stained for 15 min on ice in PBS containing 1% FCS using the antibody combinations listed in Supplementary Table 21. Data were acquired on a Fortessa X20 analyzer (BD Biosciences) and analyzed with FlowJo10 software. Flow cytometry was performed on samples from 8 WT, 9 Tg, and 5 Hum mice for the main cohort. To study *Dnm2* heterozygous mice, 6 WT, and 4 *Dnm2*^+/-^ mice were used for the analyses.

### Cytospin immunofluorescence

BM cells were cytospined for 5 min at 800 rpm and fixed with 3% PFA and 0.1% Tween-20 in PBS for 20 min at 4°C. After four washes with cold PBS, cells were blocked 1hr at 4°C with 1% FCS in PBS, then incubated overnight at 4°C with rabbit antibodies directed against human DNM2 (Thermo Fisher Scientific, #PA5-19800) or mouse DNM2 (Homemade 2865) diluted 1:100 in 1% FCS in PBS. Secondary staining was performed 1 h at room temperature (RT) with an Alexa Fluor 488-conjugated anti-rabbit IgG (H+L) antibody (Jackon ImmunoResearch #111-546-144, PA, USA) diluted 1:1000 in 1% FCS in PBS. Slides were mounted with Fluoromount containing DAPI (Invitrogen, #00-4959-52, MA, USA) and covered with 18 x 8 mm coverslips. Cytospins were imaged using a Zeiss AxioObserver Z1 time-lapse microscope.

### Transmission electron microscopy (TEM)

Whole blood samples were fixed by dilution to five times the initial volume in a fixative solution containing 4% paraformaldehyde and 0.5% glutaraldehyde. The samples were then post-fixed, pre-contrasted, dehydrated through a graded ethanol series (70%-100%), and embedded in Epon resin, as previously described^48^. Ultrathin sections (100 nm) were observed using a Jeol 2100-Plus TEM operated at 120 kV. Ten images per sample were acquired from 3 different mice per genotype at ×1500 magnification.

### Scanning electron microscopy (SEM)

Whole blood samples were fixed as described for TEM, diluted 1 :10, and 100 µL volume of the suspension was applied on poly-L-lysine-coated coverslips (Sigma-Aldrich). The coverslips were then dehydrated and dried with hexamethyldisilazane (HDMS, Merck). The coverslips were mounted on a sample holder and coated with platinum-palladium. Samples were imaged at 5kV with a Helios NanoLab DualBeam microscope (FEI). For quantification, five images per sample were acquired from 3 different mice per genotype at ×2000 magnification.

### Cell culture and lysate preparation

Hematopoietic progenitor cells (HPC-7) were cultured in an IMDM medium (Thermo Fisher Scientific, #21980) + L-Glutamine 4mM + 25mM HEPES (Life Technologies, #21980-032, CA, US) + 5% FCS + MTG 1.5x10E4 M (Sigma-Aldrich, #M6145) + Penicillin 100 UI/mL + Streptomycin 100 µg/mL. HPC-7 cells were growth in a T175 flask with media supplemented with mSCF (100ng/mL; Peprotech, #250-03, MA, US) and mIL-6 (100g/mL; Peprotech, #216-16, MA, US) until confluency was reached. Cells were transferred in a conical tube and centrifuged 5min at 4°C at 1500 rpm. The pellet was resuspended in lysis buffer (50 mM Hepes-KOH pH7.5 + 150 mM NaCl + 1% Triton X-100 + 5 mM TCEP + 10% glycerol + 1 tablet of EDTA-free Protease Inhibitor Cocktail for a final volume of 10 mL), sonicated 4x20sec, and centrifuged 20min at 4°C at 12000. Supernatant was aliquoted, complemented with 10% glycerol, and snap freeze in liquid nitrogen.

### Native holdup sample preparation

To measure binding affinities from cell extracts, we use the native holdup (nHU) assay^49^. Cytiva streptavidin high performance resin (Fisher Scientific, #15891838, MA, US) was prepared according to the manufacturer’s instructions. For saturating streptavidin resin with biotinylated peptides or biotin, 60μl of streptavidin resin was mixed with biotin or peptide at 60 μM concentration in 6 resin volume for 60 min. The following peptides synthesized by ProteoGenix (Schiltigheim, France) with a purity >98% were used as baits for the nHU experiment (underlined letters indicate aa discrepancies between human H1 & H2 and mouse M1 & M2 peptides): H1 (Biotin-Ahx-Q<u>SA</u>S<u>S</u>HSPTPQRRPVSS<u>I</u>HPPGRPPA), M1 (Biotin-Ahx-Q<u>NT</u>S<u>G</u>HSPTPQRRPVSS<u>V</u>HPPGRPPA), H2 (Biotin-Ahx-AN<u>S</u>D<u>L</u>F<u>P</u>APPQIPSRP<u>V</u>RIPPGI), and M2 (Biotin-Ahx-AN<u>N</u>D<u>P</u>F<u>S</u>APPQIPSRP<u>A</u>RIPPGI).

To saturate the free sites of the resin, 1 resin volume of 1mM biotin was added to the resin in a total volume of 9 times the resin volume for 5min at RT. After saturation, resins were washed four time with 10 resin volume in holdup buffer (50 mM Tris pH 7.5, 300 mM NaCl, and 0.8 mM TCEP, 0.22-μm filtered). 25 μl of saturated streptavidin resin was mixed with 500 μl of HPC-7 cell lysate (2 mg/ml) and incubated at 4°C for 2 hours with constant mixing. After incubation, the resin was separated from the supernatant by a 1min centrifugation at 800rpm. Then, the supernatant was removed by pipetting without any delay to avoid any resin contamination. The supernatant was centrifuged an additional time to clarify it further, removing any possible resin contamination and analyzed using an Exploris mass spectrometer.

Each biotinylated peptide condition (H1, H2, M1, M2) include biological duplicates and technical triplicates (total *n* = 6), whereas biotin control includes three biological (BIOT1, BIOT2, BIOT4) samples and technical triplicates (total *n* = 9). Mean intensities were used for fold change (FC) calculations. Protein mixtures were precipitated in trichloroacetic acid (Sigma-Aldrich) overnight at 4°C, pellets were washed twice with 1 mL cold acetone, dried and dissolved in 2 M urea (Sigma-Aldrich) in 0.1 mM Tris-HCl pH 8.5 for reduction (5 mM TCEP, 30 min.), and alkylation (10 mM iodoacetamide, 30 min.). Trypsin (Promega, WI, USA) digestion was carried out at 37°C and overnight. Peptide mixtures were then desalted on C18 spin-column and dried on Speed-Vacuum before LC-MS/MS analysis.

### Mass Spectrometry analysis and data processing

Samples were resuspended in 0.1% trifluoroacetic acid (Sigma-Aldrich) and then analyzed using an Ultimate 3000 nano-RSLC (Thermo Scientific) coupled in line with a quadrupole - orbitrap Exploris 480 via a nano-electrospray ionization source (Thermo Scientific) and the FAIMS pro interface. Less than 200 ng tryptic peptides were loaded on a C18 Acclaim PepMap100 pre-column (300 µm ID x 5 mm, 5 µm, 100Å, Thermo Fisher Scientific) for 2 minutes at 15 µL/min with 2% acetonitrile MS grade (Sigma-Aldrich), 0.1% formic acid (FA, Sigma-Aldrich) in H_2_O and then separated on a C18 PepMap nano-column (75 µm ID x 25 cm, 2.6 µm, 150Å, Thermo Fisher Scientific) with a 75 minutes linear gradient from 7% to 35% buffer B (A: 0.1% FA in H_2_O; B: 0.1% FA in 80% CAN) followed by a regeneration step at 90% B and a equilibration at 7% B. The total chromatography was 95 minutes at 450 nl/min and the oven set to 45°C. The mass spectrometer was operated in positive ionization mode in Data-Independent Acquisition (DIA) for one FAIMS compensation voltages (CV = -45V). The DIA MS parameters were set to: [1] MS: 380-985 m/z for precursor scan range; 60,000 resolution; 3E6 AGC target; 25 ms Maximum Injection Time (MIT); 60 scans events with 10 m/z windows and 1 m/z overlap. [2] HCD-MS/MS: 28% HCD collision energies; 10 m/z isolation window; 145-1450 m/z scan range; 15,000 resolution; 2E6 AGC target; 40 ms MIT; 30 loop count. Unassigned and single charged states were rejected. The exclusion duration was set for 30s with mass width was ± 10 ppm.

Proteins were identified with DIA-NN 1.9.1 software^50^ and *Mus Musculus* spectral library was generated from UniProt database (*Mus Musculus* reviewed, release 2024_04_03 with 21709 entries). Precursor and fragment mass tolerances were set at 10 ppm and 0.02 Da respectively, and up to 2 missed cleavages were allowed. Oxidation (M) and Acetyl (Protein N-term) were set as variable modification, and Carbamidomethylation (C) as fixed modification. Peptides were filtered with a false discovery rate (FDR) at 1%, rank 1. Proteins were quantified with a minimum of 1 unique peptide based on the XIC (sum of the Extracted Ion Chromatogram). The quantification values were exported in Perseus for statistical analysis involving a log[2] transform, imputation, normalization^51^. The proteomics data are available via the ProteomeXchange Consortium with the accession number PXD073642.

### Statistical Analysis

All statistical analyses and graphs were conducted using *GraphPad Prism* v9.5.1. Data normality was assessed using the Shapiro–Wilk test. One-way ANOVA with Tukey’s post-hoc test or Student’s *t*-test were performed to evaluate genotype effects; when variances between two groups were unequal, Welch’s *t*-test was used instead. For non-normally distributed data, Kruskal–Wallis test followed by uncorrected Dunn’s post-hoc test or Mann-Whitney test was applied. Two-way ANOVA with Tukey’s or Sídák’s post-hoc test was used to assess the effects of genotypes and genes. All graphs depict individual data points along with mean ± standard deviation (SD). In the case of native holdup mass spectrometry (nHU-MS) analyses, for each detected protein, a *P* value was calculated on the basis of the measured intensities of samples (*n* = 6) and controls (*n* = 9) using a two-tailed unpaired Student’s *t* test. For each nHU-MS experiment series, a hyperbolic binding threshold was calculated considering both measured intensities and the general distribution of the entire dataset, similarly as described in other works^52^. This threshold was calculated as y=y0+(−c/(x+x0)) where *y* is the *P* value threshold at fold change of *x*, *c* is a curvature parameter empirically fixed at 1, *y*_0_ is the minimal *P* value threshold, and *x*_0_ is the minimal fold change value for any given dataset. The minimal *P* value was defined at 1.3 −log_10_(*P*), and thus, there is at least 95% probability that any identified interaction partners are true interaction partners. The minimal FC value cutoff was set at 1 σ and was determined by measuring the width of the normal distribution of all measured fold changes in a given experiment. Note that this threshold can only be interpreted for interaction partners with fold change values of −*x > x*_0_.

## Study Approval

Animal care and experimentation were conducted in compliance with French and European regulations and received approval from the institutional ethics committee (project number APAFIS #58085-2025102211406712).

## Acknowledgments

We would like to thanks Dynacure and the Mouse Clinical Institute (ICS, Illkirch) for providing the *DNM2*-transgenic animals and Prof. Leif Carlsson (University of Umeå, Sweden) for providing the HPC7 cell line. We are grateful to Mohammed Selloum for his guidance on whole blood and plasma analysis, and to Aurélie Auburtin and Deborah Bitz for processing whole blood and plasma samples. We thank Hugues Jacobs for his assistance with histology interpretation, Pascal Eberling for peptide design advices, Erwan Grandgirard and the Imaging Center of the IGBMC for photonic imaging assistance, Charlotte Shmuck for her help with the nHU experiments, Bastien Morlet and the IGBMC Proteomics Facility for performing the nHU-MS experiments and data processing. We are grateful to Jean-Yves Rinckel and Fabienne Proamer for their assistance with SEM and TEM sample preparation. This work was funded by the Interdisciplinary Thematic Institute (ITI) IMCBio, as part of the ITI 2021–2028 program of the University of Strasbourg, CNRS and INSERM, was supported by IdEx Unistra (grant no. ANR-10-IDEX-0002) and the SFRI-STRATUS project (ANR-20-SFRI-0012) and EUR IMCBio (ANR-17-EURE-0023) under the framework of the France 2030 Program, and by Agence Nationale de la Recherche (Fluoproline, ANR-20-CE11-0025; CMT-GM, ANR-24-CE17-7167-01).

## Authors contribution

J.L. conceived and supervised the study and secured funding. T.A. conducted genetic crosses, collected samples, and performed molecular, cellular, and histological analyses. Pe.K. carried out flow cytometry analyses, cytospin preparations and immunostainings. C.M and A.E. performed electron microscopy analyses. C.K. assisted with genetic crosses and genotyping. Ph.K. conducted sequence alignment of DNM2 PRD domains. A.E., S.C. and Ph.K. provided project guidance. T.A. and J.L. wrote the manuscript.

---

**Supplementary Figure 1.**
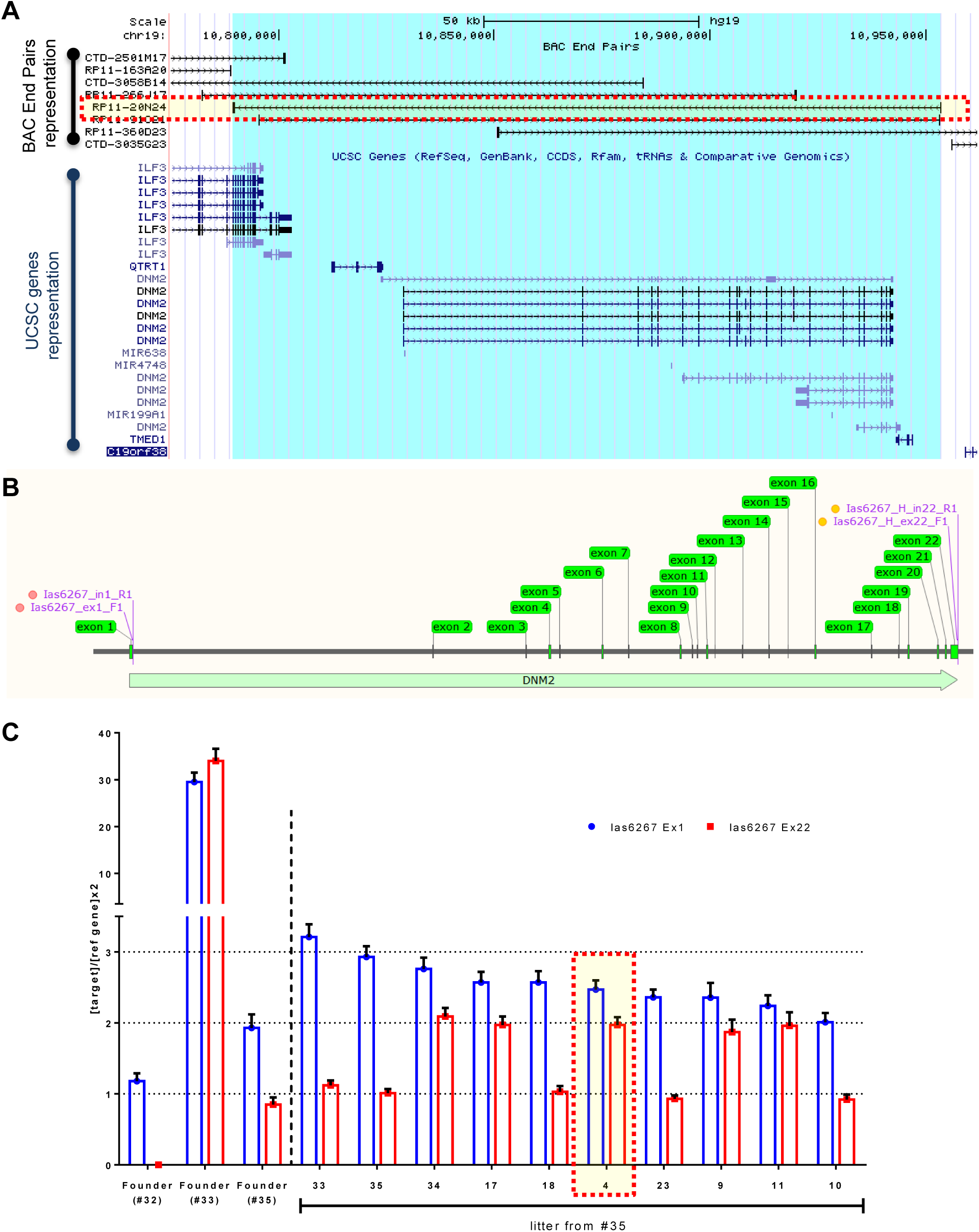
Validation of the human *DNM2* transgenic mouse. (**A**) Map of the corresponding human region on GRCh37/hg19 assembly, with the BAC sequence highlighted in blue. The human bacterial artificial chromosome (BAC) RP11-20N24 (highlighted by a dashed square) was injected in C57BL/6N embryos by DNA pronuclear injection. (**B**) Droplet Digital PCR (ddPCR) genotyping strategy. Two assays targeting exons 1 and 22 of *DNM2* were used; primer locations are indicated in purple. (**C**) ddPCR results. Founder 35 was selected, and among its offspring, animal 4 (highlighted by a dashed square), carrying two full copies and one partial copy of the BAC integrated on the Y chromosome, was selected to establish the line.

**Supplementary Figure 2.**
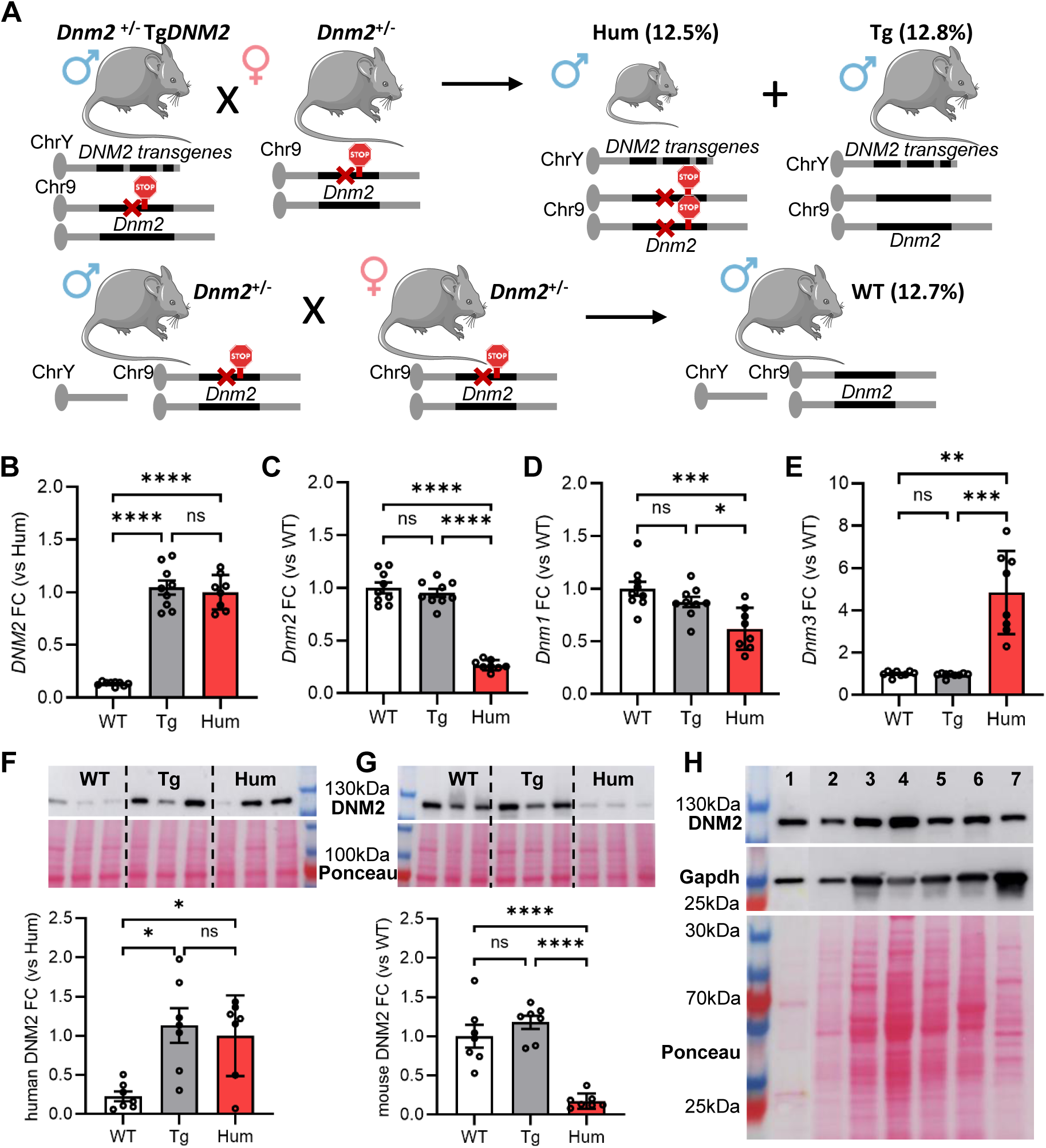
Generation and validation of the mouse models used for the study. (**A**) Mouse mating strategy used to generate mouse cohorts and ratios per genotype observed. 2-full copies of a bacterial artificial chromosome carrying the complete human *DNM2* gene were integrated on the Y chromosome. As a consequence, only males were used for this study. Mendelian ratios were obtained for wild-type (WT), transgenic (Tg), and humanized (Hum) males. (**B**-**E**) RNA fold change (FC) measurements by RT-qPCR for dynamin members using quadriceps muscles from P12 animals as samples. (**F, G**) Western blots and protein FC results using quadriceps muscles from P12 animals as samples and ponceau staining for normalization. (**H**) Western blot using humanized samples shows homogenous transgene expression across organs. Lines correspond to peripheral blood (1), spleen (2), heart (3), liver (4), kidney (5), whole brain (6), quadriceps muscle (7). Data are mean ± SD, ns: not significant, \**P* < 0.05, \*\**P* < 0.01, \*\*\**P* < 0.001, \*\*\*\**P* < 0.0001. (**B**-**D**, **G**) One-way ANOVA and Tukey’s post-hoc tests. (**E**, **F**) Kruskal-Wallis and Dunn’s post-hoc tests.

**Supplementary Figure 3.**
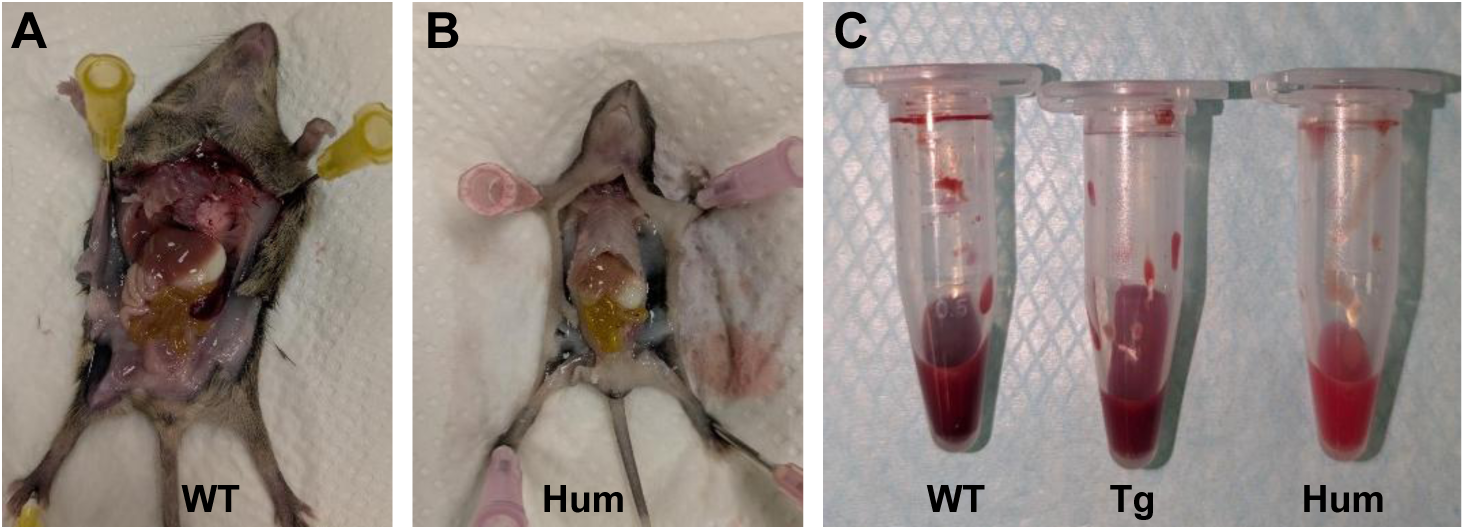
Autopsy reveals pale-colored internal organs and peripheral blood in humanized mice. (**A**, **B**) Representative dissection images of WT and Hum animals. (**C**) Blood samples obtained from WT, Tg, and Hum animals.

**Supplementary Figure 4.**
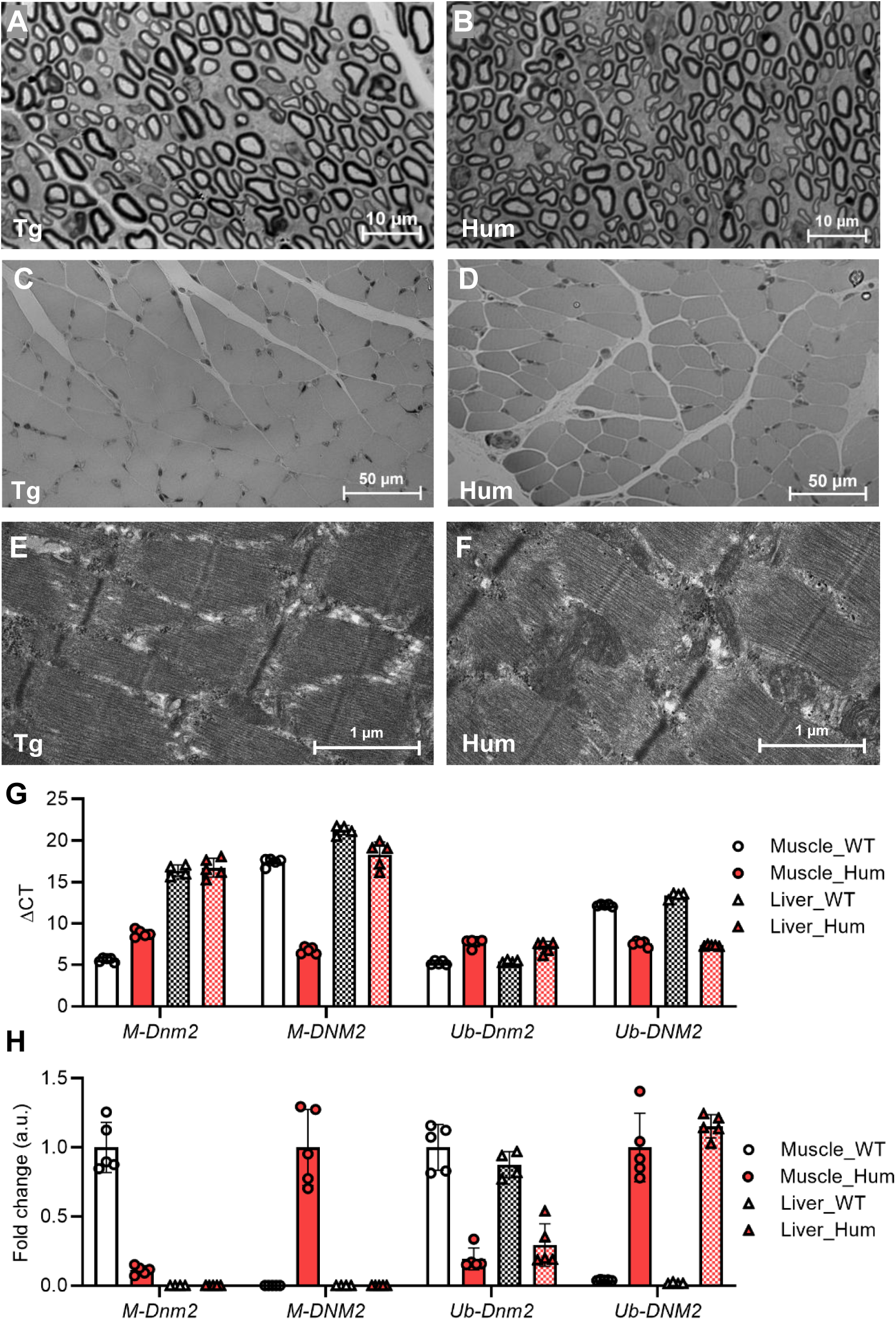
Preservation of sciatic nerve and muscle architecture in humanized and transgenic mice. (**A**, **B**) Semi-thin coronal sections of sciatic nerves. (**C**, **D**) Coronal sections of gastrocnemius muscles. (**E**, **F**) Ultrastructure analysis of gastrocnemius muscle on ultra-thin coronal sections. (**G**, **H**) Quantification of muscle-specific and ubiquitous *Dnm2*/*DNM2* isoforms expression in quadriceps and liver. ΔCT values from qPCR analysis are shown for the mouse muscle isoform (*M-Dnm2*), the human muscle isoform (*M-DNM2*), the mouse ubiquitous isoform (*Ub-Dnm2*), and the human ubiquitous isoform (*Ub-DNM2*). Higher ΔCT values correspond to lower transcript abundance. RNA fold change (FC) measurements are depicted in arbitrary units (Muscle_WT mean =1 for mouse isoforms and Muscle_Hum mean =1 for human isoforms). Data are mean ± SD.

**Supplementary Figure 5.**
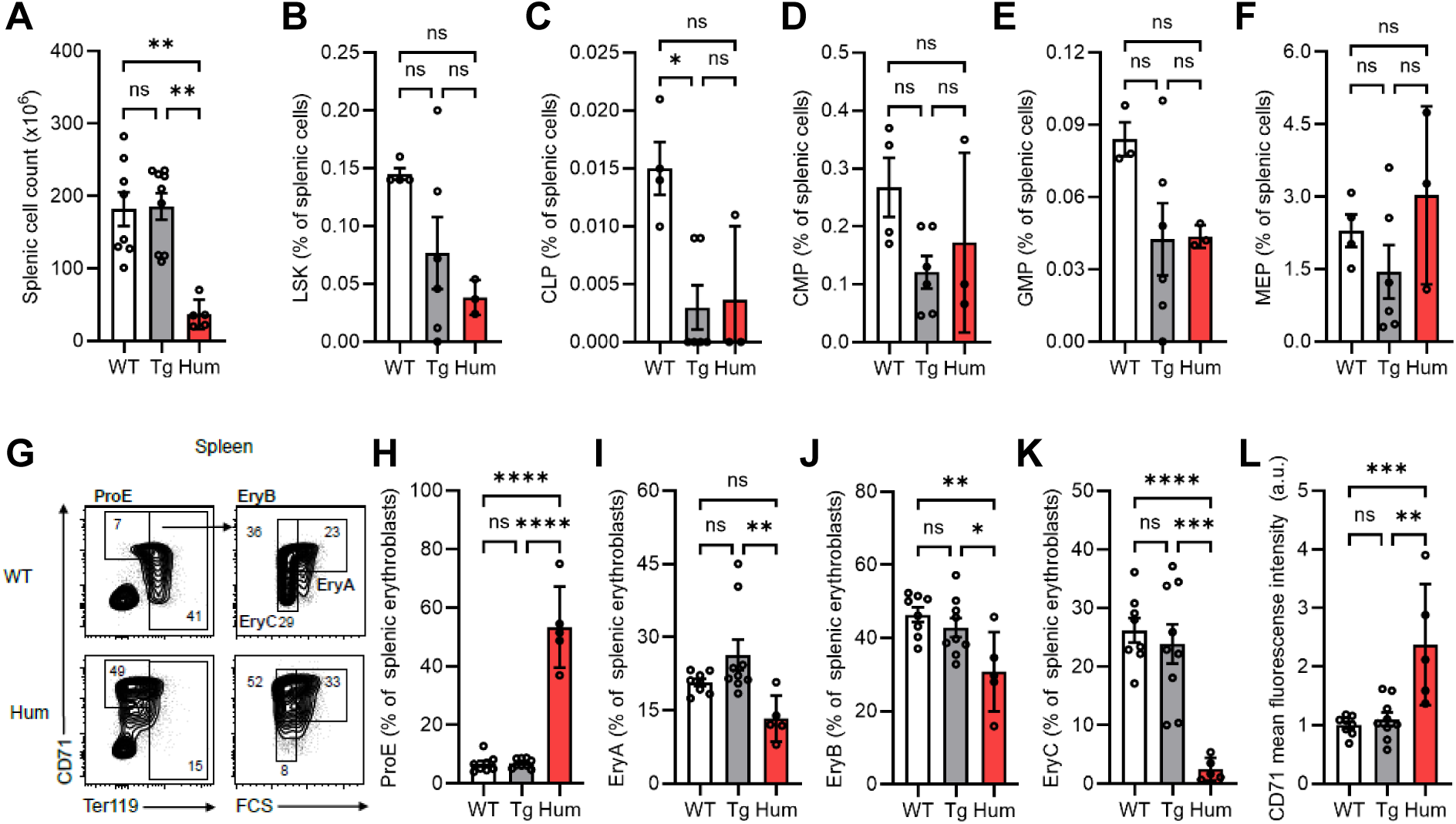
Humanized mice show impaired erythroblast maturation in the spleen. (**A**) Cell count derived from total spleens. (**B**-**F**) LSK/LK hematopoietic progenitors (% cells). LSK: Lin^-^Sca1^+^c-Kit^+^ hematopoietic stem and progenitor cells. CLP: Common lymphoid progenitors. CMP: Common myeloid progenitors. GMP: Granulo-monocyte progenitors. MEP: Megakaryo-erythrocyte progenitors. (**G**) Representative flow cytometry plots using CD71 and Ter119 expressions with the indicative percentage of cells. (**H**-**K**) Erythroblast populations (% erythroblasts). ProE: proerythroblasts. EryA: basophilic erythroblasts. EryB: polychromatophilic erythroblasts. EryC: orthochromatic erythroblasts. (**L**) CD71 mean fluorescence intensity in erythroblasts (arbitrary units). Data are mean ± SD, ns: not significant, \**P* < 0.05, \*\**P* < 0.01, \*\*\**P* < 0.001, \*\*\*\**P* < 0.0001. (**D**-**F**, **H**, **J**-**L**) One-way ANOVA + Tukey’s post-hoc; (**A**-**C**, **I**) Kruskal-Wallis + Dunn’s post-hoc.

**Supplementary Figure 6.**
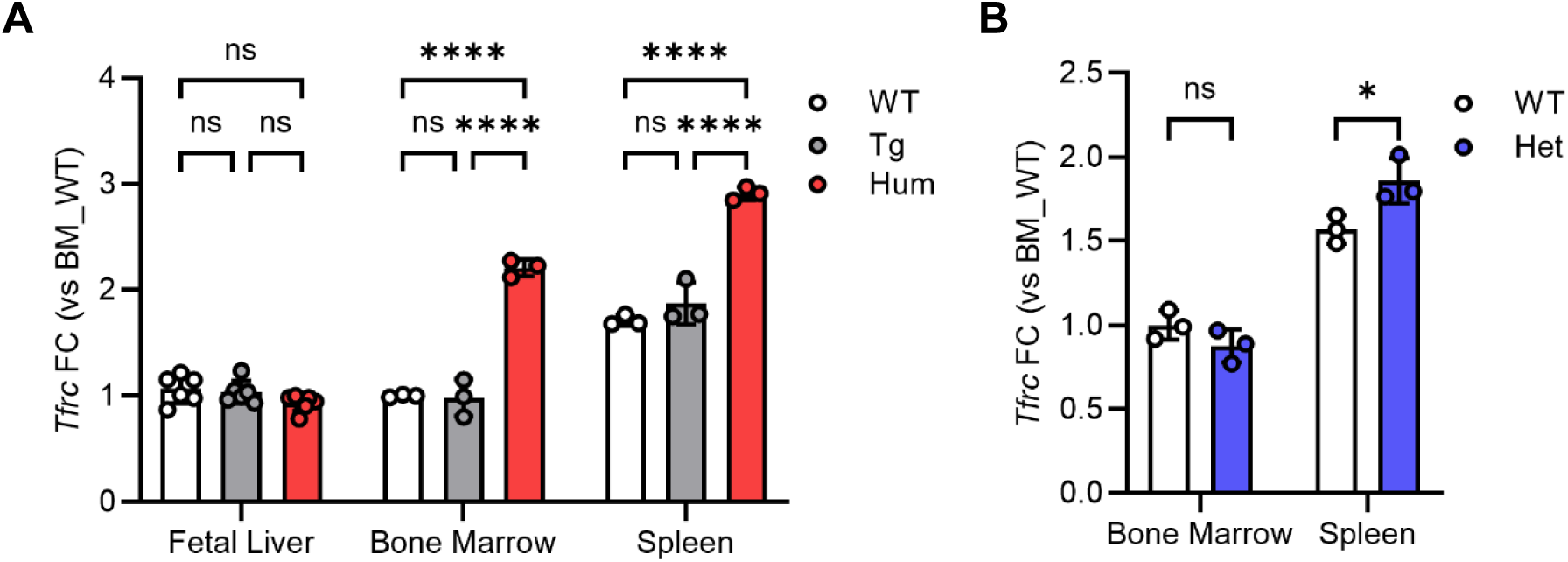
The *Tfrc* gene, encoding the transferrin receptor CD71, is dysregulated across multiple tissues in *DNM2-*Hum and *Dnm2*-Het mice. (**A**) *Tfrc* RNA fold-change (FC) values in E14.5 fetal liver and in P12 bone marrow (BM), spleen, and quadriceps. (**B**) *Tfrc* RNA fold-change (FC) values in P12 BM and spleen. Data are mean ± SD in arbitrary units (BM_WT mean =1), ns: not significant, \**P* < 0.05, \*\*\**P* < 0.001, \*\*\*\**P* < 0.0001. (**A**) Two-way ANOVA + Tukey’s post-hoc. (**B**) Two-way ANOVA + Sídák’s post-hoc.

**Supplementary Figure 7.**
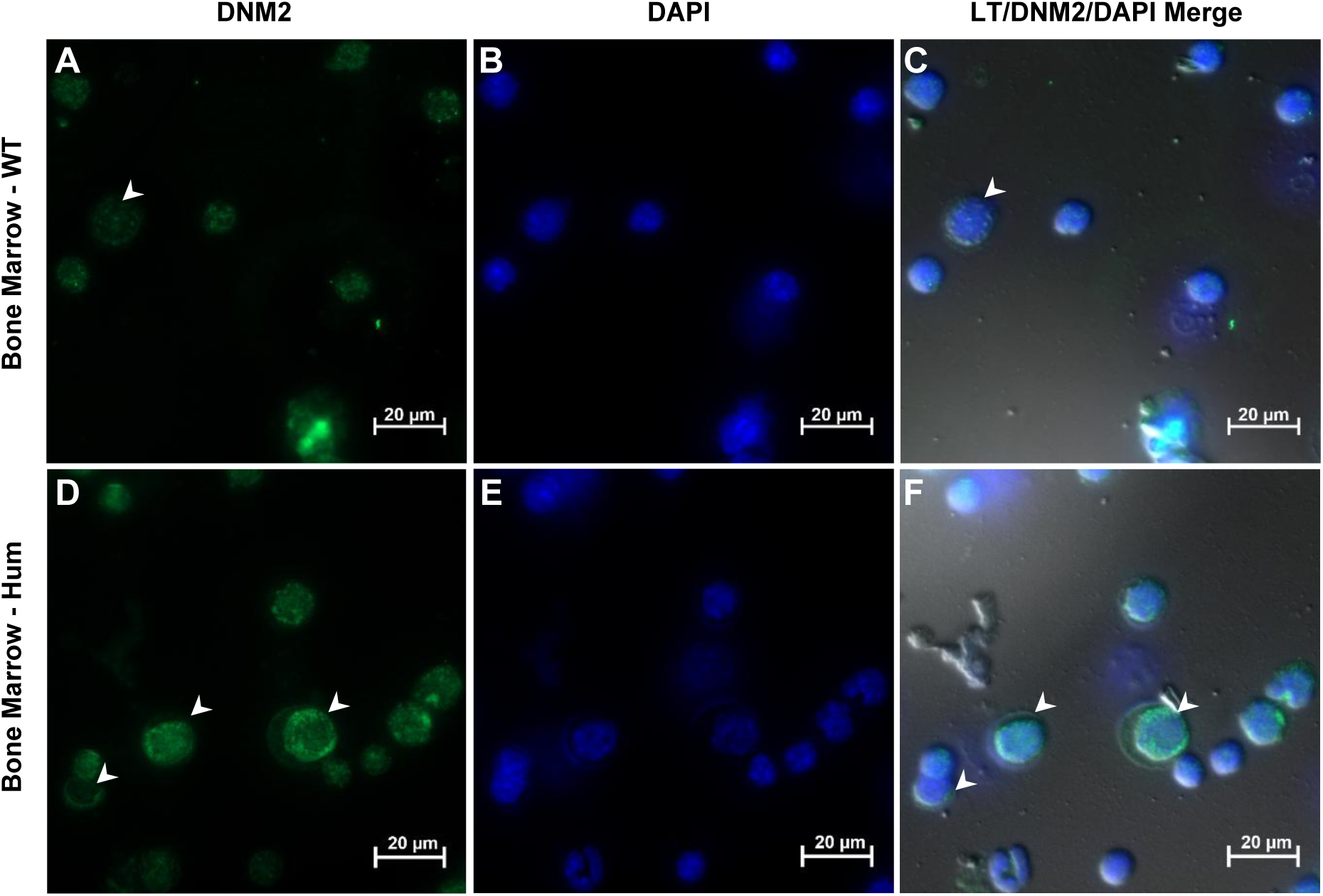
Time-lapse microscope immunofluorescence of bone marrow cytospin preparations shows perinuclear and cytoplasmic localization of human and mouse DNM2 proteins. (**A**-**C**) WT samples. (**D**-**F**) Hum samples. (**A**, **D**) DNM2 immunostaining; (**B**, **E**) DAPI nuclear staining; (**C**, **F**) Merged light transmission (LT)/DNM2/DAPI images. The human DNM2 antibody was used for Hum samples, whereas the mouse DNM2 antibody was used for WT samples. White arrowheads highlight cells displaying clear perinuclear and cytoplasmic DNM2 localization. Note that in nucleated hematopoietic cells, most cell volume is occupied by the nucleus.

**Supplementary Figure 8.**
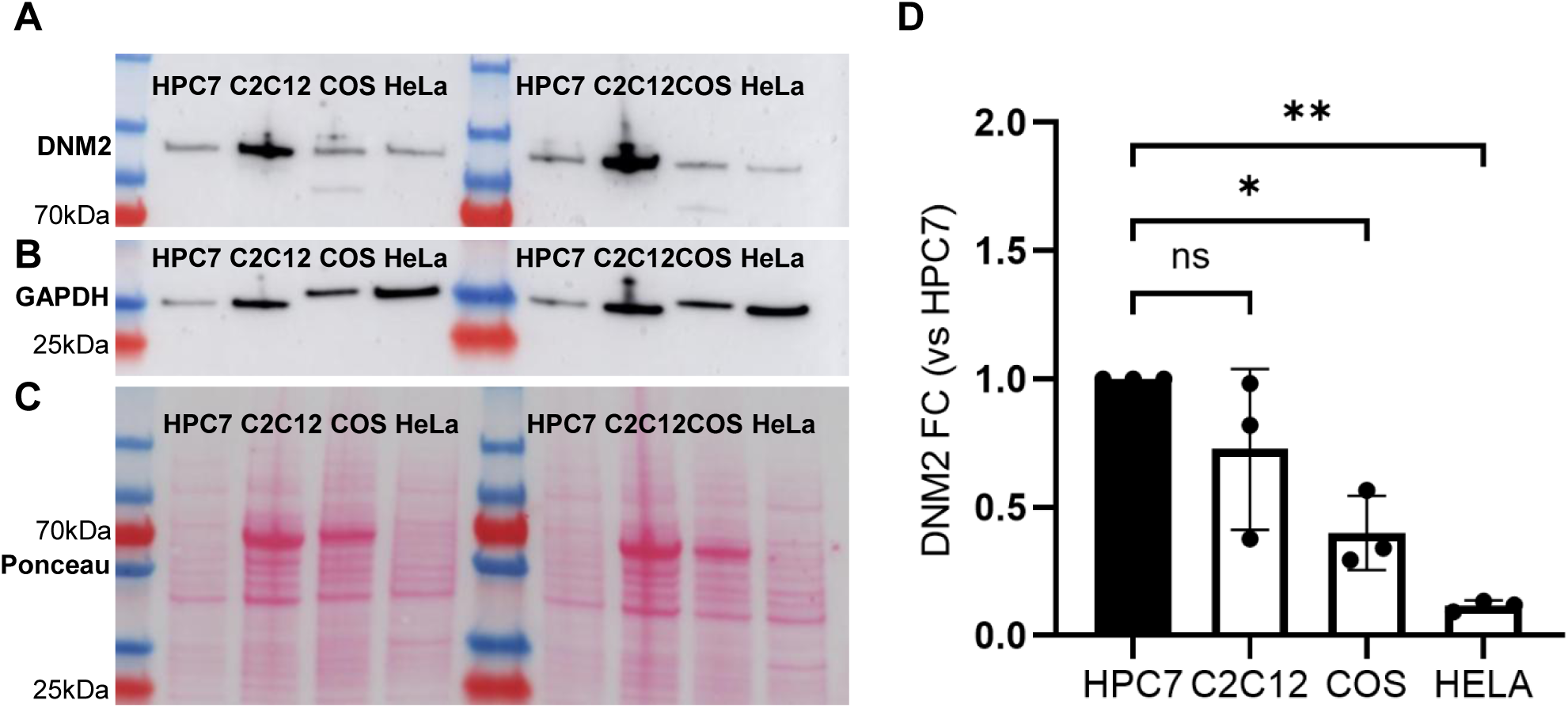
Western blot analysis of DNM2 protein across different cell types showing the higher expression in HPC7 cells. (**A**, **B**) Western blot membranes using a primary antibody against DNM2 and GAPDH. (**C**) Ponceau staining. (**D**) DNM2 protein fold change (FC) results using GAPDH for normalization. HPC7: mouse hematopoietic progenitors cell line. C2C12: mouse myoblast cell line. COS: monkey fibroblast-like cell line. HeLa: human cervical adenocarcinoma cell line. Data are mean ± SD, ns: not significant, \**P* < 0.05, \*\**P* < 0.01. One-way ANOVA and Tukey’s post-hoc tests.

**Supplementary Figure 9.**
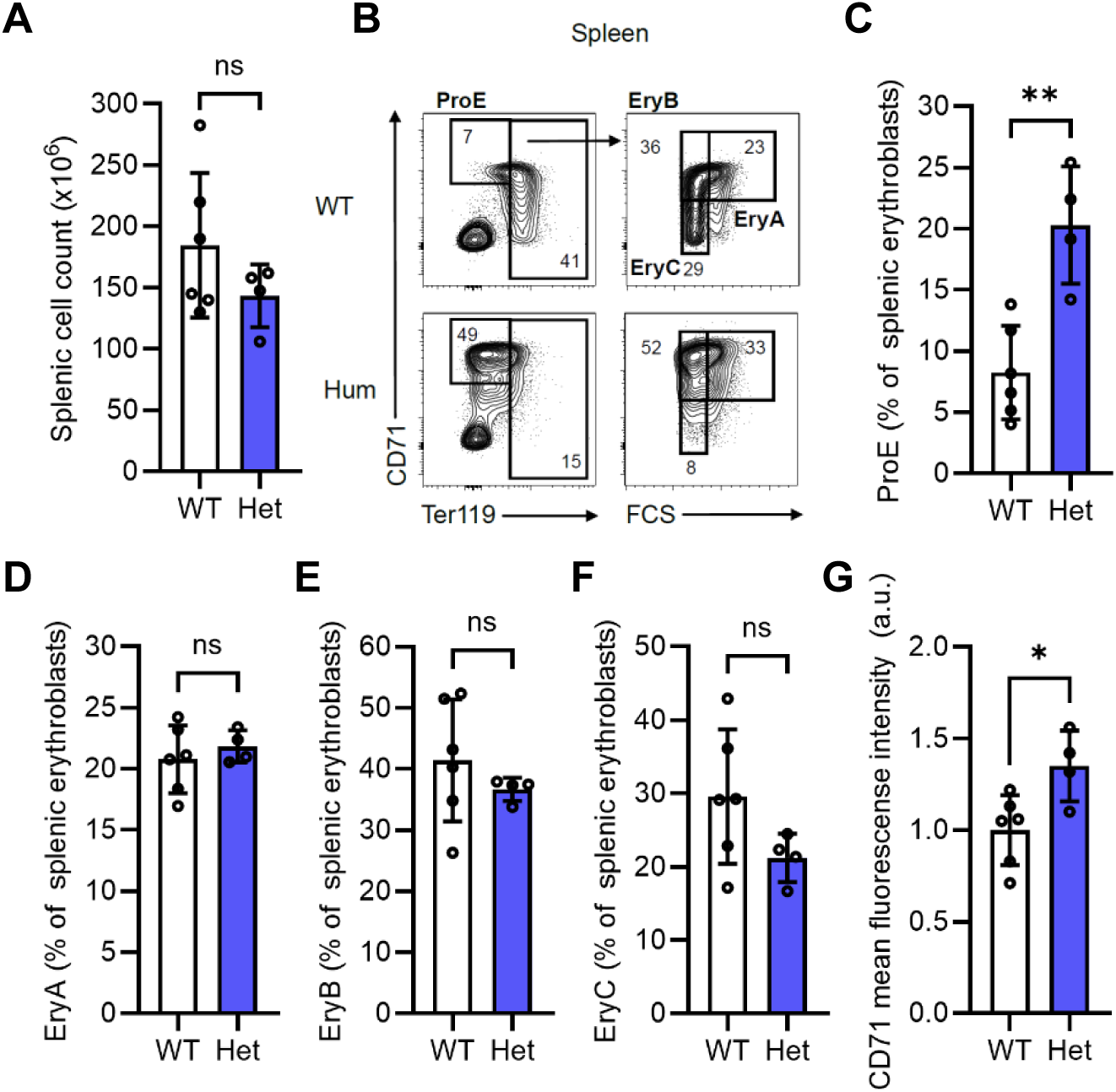
*Dnm2* haploinsufficiency results in an accumulation of proerythroblast and increased CD71 cell surface signal in the spleen. (**A**) Cell counts derived from total spleens (x10^6^ cells). (**B**) Representative flow cytometry plots using CD71 and Ter119 expressions with the indicative percentage of cells. (**C**-**F**) Erythroblast populations (% erythroblasts). (**G**) CD71 cell surface mean fluorescence intensity in erythroblasts (arbitrary units). Data are mean ± SD, ns: not significant, \**P* < 0.05, \*\**P* < 0.01. (**A**, **C**, **D, F**, **G**) Student’s *t*-test; (**E**) Mann-Whithey test.

**Supplementary Figure 10.**
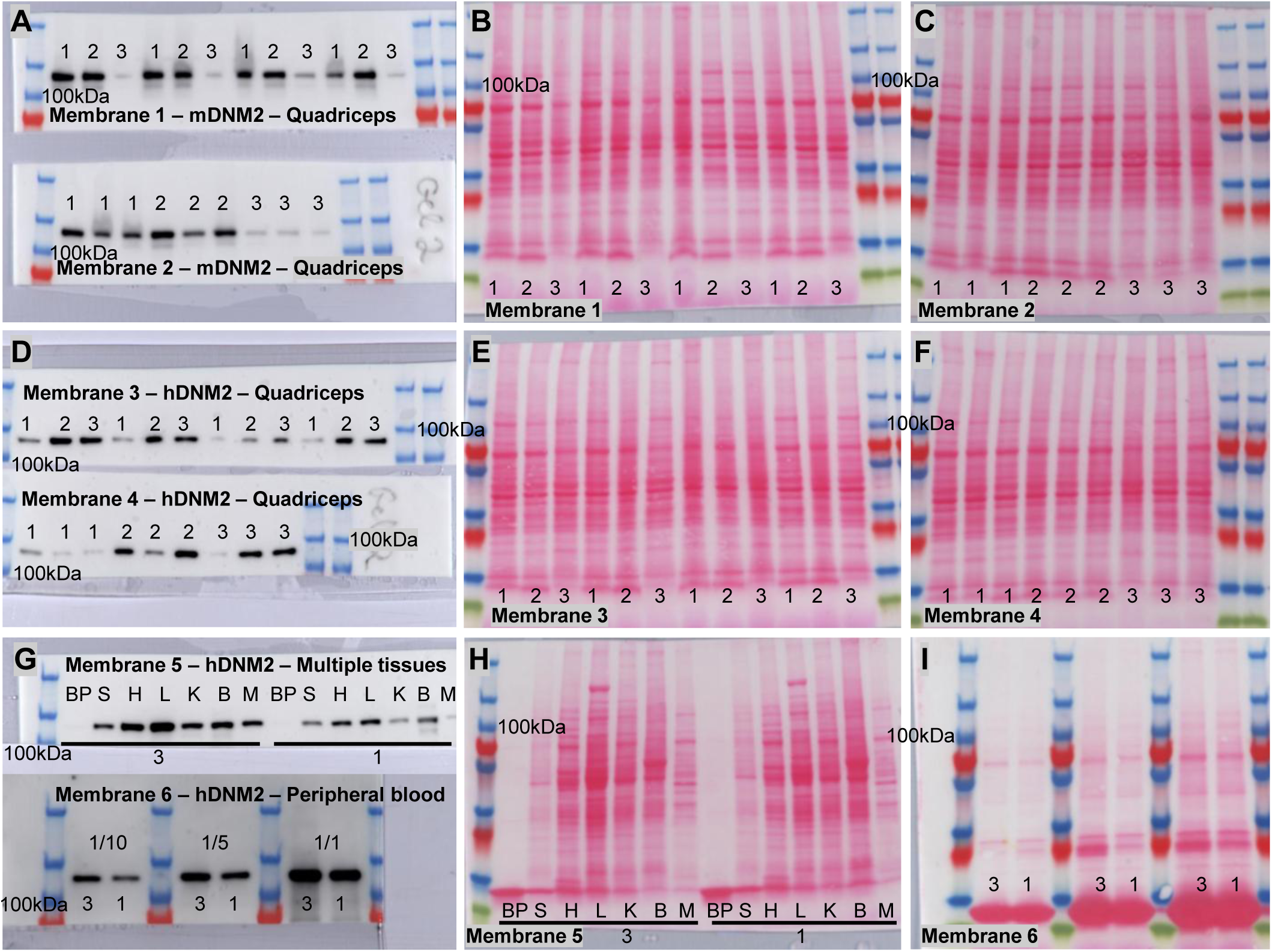
Western blotting membranes used for protein quantification. (**A**) Blots using a primary antibody against mouse DNM2. (**D**, **G**) Blots using a primary antibody against human DNM2. (**B**, **C**, **E**, **F**, **H**, **I**) Ponceau stainings used for protein normalization. (**A**-**F**) Quadriceps samples. 1: WT samples; 2: Tg samples; 3: Hum samples. (**G**-**I**) Multiple organ samples. 1: WT samples; 3: Hum samples. BP: peripheral blood pellet; S: spleen; H: heart; L: liver; B: total brain. In membrane 5, blood pellet samples were used following 5000G centrifugation. In membrane 6, blood samples were used at different dilutions.

